# E-proteins set the threshold for optimal TCF1 expression during αβ T cell development

**DOI:** 10.1101/2023.11.06.565822

**Authors:** Anjali Verma, Bridget Aylward, Fei Ma, Cheryl A. Sherman, Laura Chopp, Susan Shinton, Roshni Roy, Shawn Fahl, Alejandra Contreras, Byron Koenitzer, Parirokh Awasthi, Krystyna Mazan-Mamczarz, Supriyo De, Noah Ollikainen, Xiang Qiu, Remy Bosselut, Ranjan Sen, David L. Wiest, Jyoti Misra Sen

**Affiliations:** Laboratory of Clinical Investigation National Institute on Aging 251 Bayview Boulevard Baltimore, MD 21224; Nuclear Dynamics and Cancer Program Fox Chase Cancer Center, 333 Cottman Avenue Philadelphia, PA 19111; Laboratory of Molecular Biology and Immunology National Institute on Aging, 251 Bayview Boulevard Baltimore, MD 21224; Laboratory of Immune Cell Biology Center for Cancer Research, National Cancer Institute National Institutes of Health Bethesda, MD 20892; Mouse Modeling & Cryopreservation (MMC) The NCI Mouse Repository Laboratory Animal Sciences Program National Cancer Institute Frederick, MD 21702; Laboratory of Genetics & Genomics National Institute on Aging 251 Bayview Boulevard Baltimore, MD 21224; Immunology Program Department of Medicine Johns Hopkins School of Medicine Baltimore, MD 21224

**Author notes:** Equal contribution (Anjali Verma, Bridget Aylward, Fei Ma, Cheryl A. Sherman).

## Abstract

Expression of T Cell Factor-1 (TCF1), encoded by *Tcf7,* regulates lineage fate decisions during T cell development. Here we demonstrate that E-proteins control the threshold of TCF1 expression required for development of T cells. E-proteins bind to five elements (EPEs) in the *Tcf7* locus. The third element, EPE3, interacts directly with *Tcf7* promoter in Hi-ChIP analyses, suggesting it is an active enhancer. CRISPR-ablation of EPE3 reduces TCF1 protein expression in precursor thymocytes by 2-fold and dramatically impairs development of αβ and γδ T cells. Single cell gene expression analysis identified differentiation blocks at multiple CD4^-^CD8^-^ stages and subsequent transition to CD4^+^CD8^+^ stage. These data identify E-proteins and EPE3 as critical for the optimal TCF1 expression required for T cell development.

## Introduction

T cell factor-1 (TCF1), encoded by *Tcf7* gene, plays an essential role in the development of both conventional and innate T cells ^1, 2, 3, 4, 5, 6, 7^. TCF1 is a necessary and sufficient effector of Notch signals that impose the T lineage fate on multipotent thymic CD4^-^CD8^-^ double negative (DN) progenitors ^8, 9, 10^. Specifically, Notch signals initiate transcription of *Tcf7* in CD44^+^CD25^-^ DN1 cells ^8, 10, 11, 12^, leading to induction of Bcl11b and T lineage commitment in CD44^-^CD25^+^ DN2 cells ^13, 14, 15^. TCF1 promotes the development of DN αβ lineage progenitors to the CD4^+^CD8^+^ double positive (DP) stage in cooperation with the E-protein HEB ^2, 5, 6^, while opposing γδ fate specification ^16^. Finally, TCF1 plays a critical role in lineage commitment of DP thymocytes to single positive (SP) cells by inducing *Zbtb7b* to promote the CD4 fate, and along with *Runx3*, silencing CD4 in CD8 SP thymocytes ^17^.

Along with its critical role in supporting the development of conventional αβ lineage T cells, TCF1 also influences innate-like T lineages including NKT cells and IL-17 producing γδ T cells ^16, 18, 19^. TCF1 promotes the development of NKT cells by prolonging the survival of DP thymocytes and enabling rearrangement of the distal Vα elements required to assemble the stereotypic T cell receptor (TCR) complex employed by NKT cells ^19^. TCF1 also plays an important role in regulating the development of innate-like IL-17 producing γδ T cells by antagonizing the function of the fate specifying transcription factor RORγt ^16, 18^.

The ability of TCF1 to influence multiple cell fates during T cell development of adaptive and innate lymphoid lineages is determined by its expression level ^20^. For example, development of conventional αβ lineage T cells requires the maintenance of high levels of expression of TCF1, while γδ lineage commitment requires TCF1 expression to be reduced ^16^. How TCF1 expression levels are controlled remains unclear. While Notch signals initiate *Tcf7* expression ^8, 10^, the extent to which Notch1 signaling regulates TCF1 expression during subsequent developmental transitions and checkpoints is not known. A recent report identified a 1kb regulatory element upstream of the *Tcf7* locus that plays a critical role in controlling TCF1 expression; however, ablation of the Notch binding sites in this element had minimal impact on TCF1 expression, suggesting other signaling axes and transcription factors play a more predominant role in setting TCF1 expression levels ^21^. We have recently demonstrated that the activity of E box DNA binding proteins play an important role in regulating TCF1 expression in γδ lineage T cells and defined one E-protein element through which the regulation occurs ^16^. However, mechanisms that maintain TCF1 expression levels during αβ T cell development and maturation remain unknown.

In this report, we systematically evaluate the role of E-proteins and the regulatory elements to which they bind in controlling TCF1 expression levels in developing thymocytes. ChIP-Seq for E-proteins E2A and HEB to define the E-protein regulome upstream of the *Tcf7* gene, revealed five E-protein bound regulatory elements (EPEs). The function of each EPE was assessed by CRISPR deletion in mice, revealing that each EPE contributes differently towards *Tcf7* regulation in γδ and conventional αβ T cell precursors. While ablation of EPE1, 3, and 5 reduced TCF1 expression in γδ T cells and affected their maturation and adoption of the IL-17 producing effector fate, ablation of 207bp encompassing EPE3 (ΔEPE3 mice) was the only EPE whose ablation reduced TCF1 expression in αβ T cells and attenuated their development. Single cell RNAseq analysis of DN progenitors from ΔEPE3 mice revealed that transcriptomic disruption began in DN2 thymocytes and persisted through the developmental block at the CD44^-^CD25^+^ (DN3) to CD44^-^CD25^-^ (DN4) transition, where pre-TCR mediated β-selection occurs ^22^. We also provide mechanistic insight into EPE3 function, since it was found to be epigenetically marked with H3K27 acetylation (H3K27ac) and to interact with the *Tcf7* promoter, which was disrupted in ΔEPE3 thymocytes. Importantly, the EPE3 deletion preserved the adjacent Notch binding site; however, ectopic expression of Notch failed to correct the reduced TCF1 expression observed in progenitors from ΔEPE3 mice. We conclude that after Notch-dependent initiation, the expression level of TCF1 is controlled by E-protein binding to a series of EPE sites, of which EPE3 determines the threshold required for optimal development of conventional αβ T cells.

## Results

### E-protein regulome in *Tcf7* regulatory region

Our previous analysis indicated that the induction of Id3 by strong TCR signals repressed the expression of TCF1 in γδ lineage precursors by repressing the activity of E box DNA binding proteins ^16^. Consequently, we analyzed our previously reported ChIP-Seq data to assess the binding of E proteins E2A and HEB near the *Tcf7* locus in DN3 progenitors ^23, 24^. E-proteins bound to 5 regions (EPE) dispersed over a 60kb genomic interval beginning in the first intron of the *Tcf7* gene (Fig. 1A and Suppl. Fig. 1). Each EPE contains 2 to 4 consensus E-protein binding motifs. In a previous study, we showed that EPE1 controlled *Tcf7* expression selectively in γδ T cells and regulated their lineage commitment and maturation ^16^. To gain a comprehensive view of the ability of all the EPEs to control *Tcf7* expression, we carried out Hi-ChIP in DN3 cells using anti-H3K27ac antibodies.

**Figure 1.**
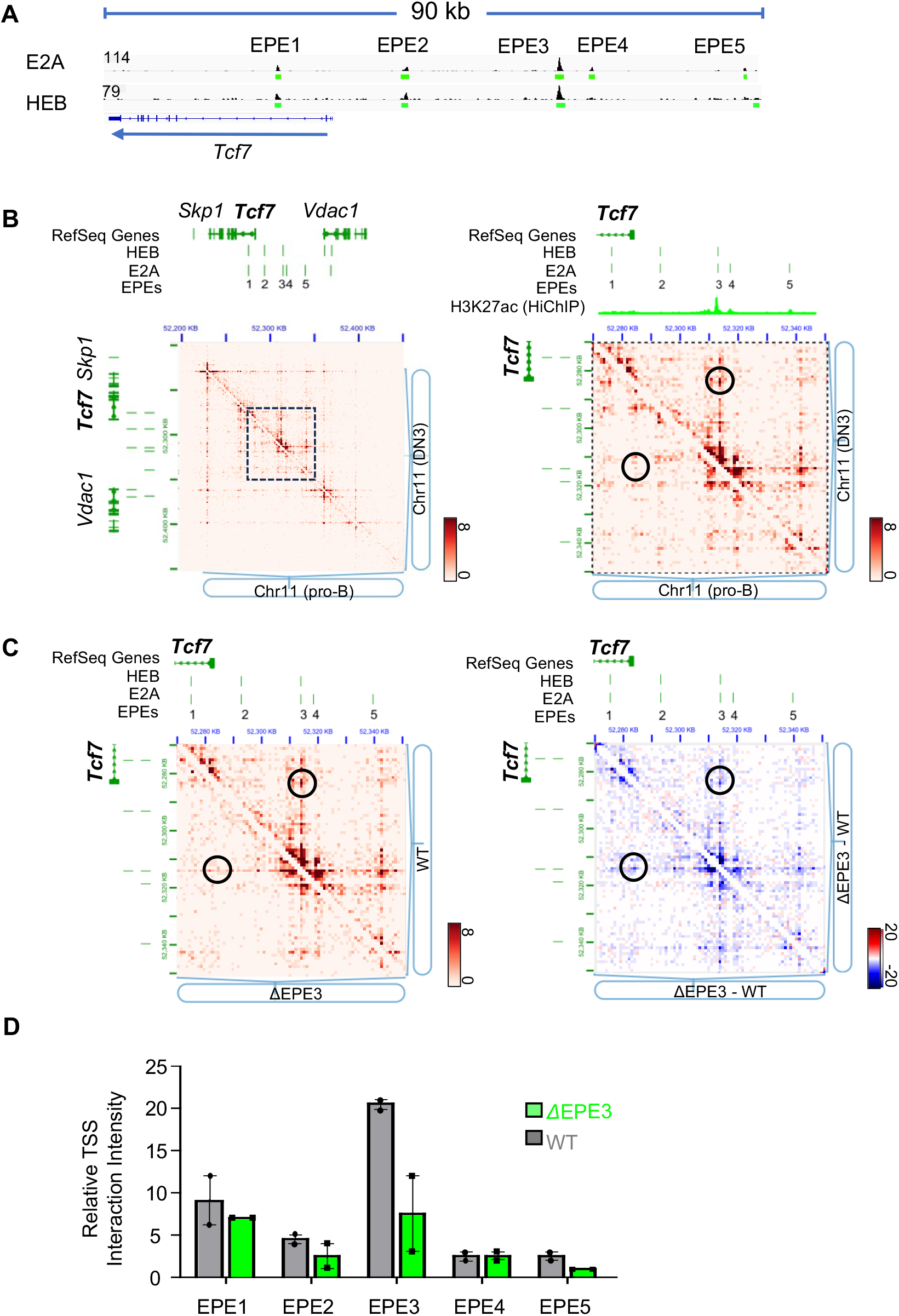
Unveiling regulatory networks at the *Tcf7* promoter by epigenetic signatures and cell type specific interactions. **(A)** IGV tracks showing E2A and HEB ChIP-seq data from fetal DN3 thymocytes. Five distinct peaks (EPE1-5) of E2A and HEB binding were identified in the vicinity of the *Tcf7* locus. **(B)** Interaction heat maps illustrating H3K27ac Hi-ChIP signals in *Rag2*^-/-^ pro-B (bottom left) and DN3 (upper right) cells, presented at 1kb resolution. The region spanning the 5 EPEs adjacent to the *Tcf7* locus is demarcated by a black dashed box, enlarged in the inset for closer examination on the right panel. ChIP-seq peaks featured in (**A)** are indicated on the top of the heatmaps. The right-side magnified map also showcases the H3K27ac peak track derived from H3K27ac Hi-ChIP data. The DN3 specific interaction is annotated by black circle. The scale bars indicate the contact frequency. **(C)** Interaction heat maps illustrating H3K27ac Hi-ChIP signals within the *Tcf7* promoter region in ΔEPE3 deletion (bottom left) and wild-type (upper right) DN3 cells, presented at 1kb resolution. The scale bars indicate the contact frequency. The difference map (ΔEPE3 minus WT) for the same region is shown on the right; regions with increased and decreased interactions are colored red and blue, respectively. The scale bar indicates the difference in contact frequency. ChIP-seq peaks featured in (**A**), are indicated on the top of the heatmaps. Interactions between EPE3 and *Tcf7* TSS are indicated by black circles. **(D)** Bar graphs displaying the relative interactions between *Tcf7* TSS and the five E2A peaks in WT and ΔEPE3 DN3, as featured in (A). These interactions are quantified within both wild-type and EPE3 mutant DN3 cells, with error bars representing the standard error of the mean (SEM) for 2 replicates.

Hi-ChIP constitutes enriching a subset of genomic DNA fragments from a Hi-C library using antibodies prior to deep sequencing. Whereas Hi-C scores for ‘all-to-all’ proximity ligations, Hi-ChIP identifies those associated with specific proteins^25^. Consequently, anti-H3K27ac Hi-ChIP identifies genomic interactions of regions that contain H3K27ac histone modification, which is associated with active promoters and enhancers^26, 27^. We found that the *Tcf7* promoter selectively interacted with EPE3 and to a lesser extent with EPE5 in DN3 cells, but not in bone marrow-derived pro-B cells lacking TCF1 expression (Fig. 1B). By contrast, long distance interactions between the *Vdac1* and *Skp1* genes were present in both DN3 and pro-B cells (Fig. 1B). These observations suggest that EPE3, which contains three putative E-protein binding sites, may play an important role in controlling *Tcf7* expression in developing thymocytes. To test this, we employed CRISPR/Cas9 technology to ablate a 207 bp region containing the three E2A/HEB binding sites in EPE3 (Suppl. Fig. 1B). Importantly, the nearby Notch binding site (NBS) was left intact to distinguish effects of Notch signaling from those of E2A/HEB binding (Suppl. Fig. 1A). Hi-ChIP analysis of DN3 thymocytes revealed that ablation of EPE3 reduced interactions between the chromatin around EPE3 and the transcription start site (TSS) of the *Tcf7* gene (Fig. 1B-D). Interestingly, interaction between EPE5 and the TSS also appear to be modestly reduced in DN3 cells in which EPE3 was ablated (Fig. 1B, quantified in 1D). Thus, the 207bp EPE3 sequence containing the 3 E-protein binding sites is required for optimal interactions with the TSS of the *Tcf7* gene.

### EPE3 deletion **r**educes levels of TCF1 expression and disrupts T cell development

Reduced interaction between EPE3 and the TSS of *Tcf7* on ΔEPE3 alleles was reflected in lower levels of TCF1 protein expression in all thymocyte subsets (Fig. 2A, B). The reduction in TCF1 expression was accompanied by markedly reduced thymic cellularity and the number of DP thymocytes, which were 4-fold lower in ΔEPE3 mice (Fig. 2C, D). Fewer DP precursors also resulted in reduced generation of SP thymocytes (Fig. 2C, D). Similar observations were made in three independent ΔEPE3 founder lines (Suppl. Fig. 2). The removal of EPE1, EPE2, and EPE4 had no substantial impact on αβ T cell development, whereas deletion of EPE5 exhibited modest effects (Suppl. Fig. 3).

**Figure 2:**
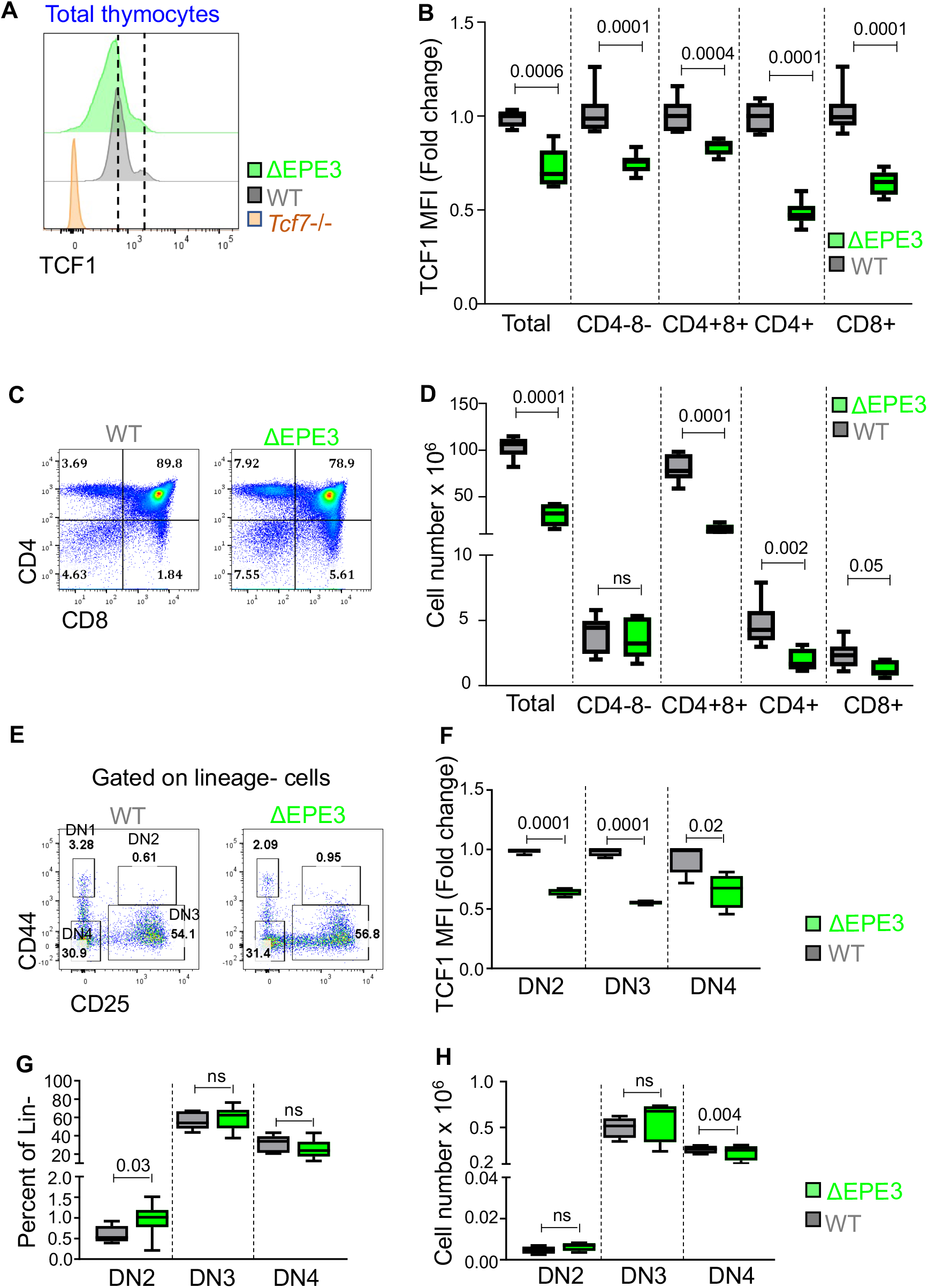
EPE3 regulates TCF1 expression in developing thymocytes. **(A)** Intracellular flow cytometry analysis of expression of TCF1 in thymocytes of WT, ΔEPE3 and *Tcf7*-/- mice. Dashed lines denote mean peak intensity for the two peaks in WT thymocytes. **(B)** Graph represents fold-change in mean fluorescent intensity (MFI) of TCF1 in the indicated ΔEPE3 thymic subset relative to their WT counterpart. WT (n=4), and ΔEPE3 (n=8) mice. **(C)** Flow cytometry analysis of thymic subsets defined by surface expression of CD4 and CD8 in thymus of WT, and ΔEPE3 mice. **(D)** Absolute number of the indicated thymic subsets: Total thymocytes, double negative (CD4- CD8-), double positive (CD4+CD8+), CD4 single positive and CD8 single positive cells. WT (n=8), and ΔEPE3 mice (n=7). **(E)** Flow cytometry analysis of DN2 (lineage-CD44+CD25+) -DN4 cells defined by surface expression of CD44 and CD25 in lineage-thymocytes from WT, and ΔEPE3 mice. **(F)** Graphic representation of the fold-change in TCF1 MFI in the indicated ΔEPE3 DN subset relative to their WT counterpart: DN2 cells (lineage-CD44+CD25+), DN3 cells (lineage-CD44- CD25+), and DN4 cells (lineage-CD44- CD25-) cells. WT (n=5), and ΔEPE3 (n=5) mice. (**G, H**) Graphical representation of the frequencies **(G)** and number **(H)** of DN2-4 cells in the thymi of WT (n=6), and ΔEPE3 mice (n=12). Results are representative of 3 or more experiments performed and p values are indicated as noted in methods.

To identify the stage at which the developmental defect in ΔEPE3 thymocytes originated, we subdivided the DN population based on changes in CD44 and CD25 expression. We found that TCF1 expression was reduced in DN2, DN3 and DN4 thymocytes (Fig. 2E, F). The proportions and cell numbers of DN2 and DN3 subsets were unaffected; however, DN4 cell numbers were modestly decreased (Fig. 2G, H). The reduction in DN4 cell numbers was consistent with the primary impairment of αβ T cell development in ΔEPE3 mice occurring in transition to the DP stage.

We previously showed that deleting EPE1 reduced TCF1 expression in γδ T cell precursors, advanced their maturation, and promoted adoption of the IL-17 producing effector fate ^16^. ΔEPE3 mice exhibited similar, but more profound, effects on γδ T cell maturation congruent with further reduction in TCF1 expression (Suppl. Fig. 4, 5). Deletion of EPE2 and 4 had no effect on γδ maturation, while deletion of EPE5 modestly promoted γδ T cell maturation and an increase in IL- 17 production (Suppl. Fig. 6). Together these data indicate that each EPE has lineage-restricted functions with EPE1 and EPE5 primarily supporting development of γδ T cells, while EPE3 regulates development of both γδ and conventional αβ T cells. It is surprising that the dramatic effects of EPE3 deletion on αβ T cell development were manifest with less than 2-fold reduction in TCF1 levels. These observations emphasize the exquisite sensitivity of particular developmental transitions to the expression level of TCF1.

### Reduced expression of TCF1 in ΔEPE3 DN thymocytes impedes transition of DN thymocytes **to the DP stage.**

Pre-TCR signaling in DN3 thymocytes promotes differentiation into DN4 and then DP thymocytes, which is accompanied by and depends upon robust proliferation ^28^. This process can be kinetically analyzed *in vitro* by single cell suspension culture of DN cells or by co-culture on monolayers of OP9-DL1 cells ^29, 30, 31^. To better define the stage at which ΔEPE3 thymocytes were developmentally impaired, we cultured purified DN4 thymocytes *in vitro* and assayed their development into DP thymocytes (Fig. 3A, B). Fewer ΔEPE3 DN4 thymocytes transitioned to the DP stage compared to WT cells, while a greater proportion remained as DN thymocytes (Fig. 3B). Similar results were observed when total DN thymocytes were co-cultured on OP9-DL1 monolayers (Fig. 3C-E), indicating that ablation of EPE3 attenuated development of DN4 thymocytes to the DP stage.

**Figure 3:**
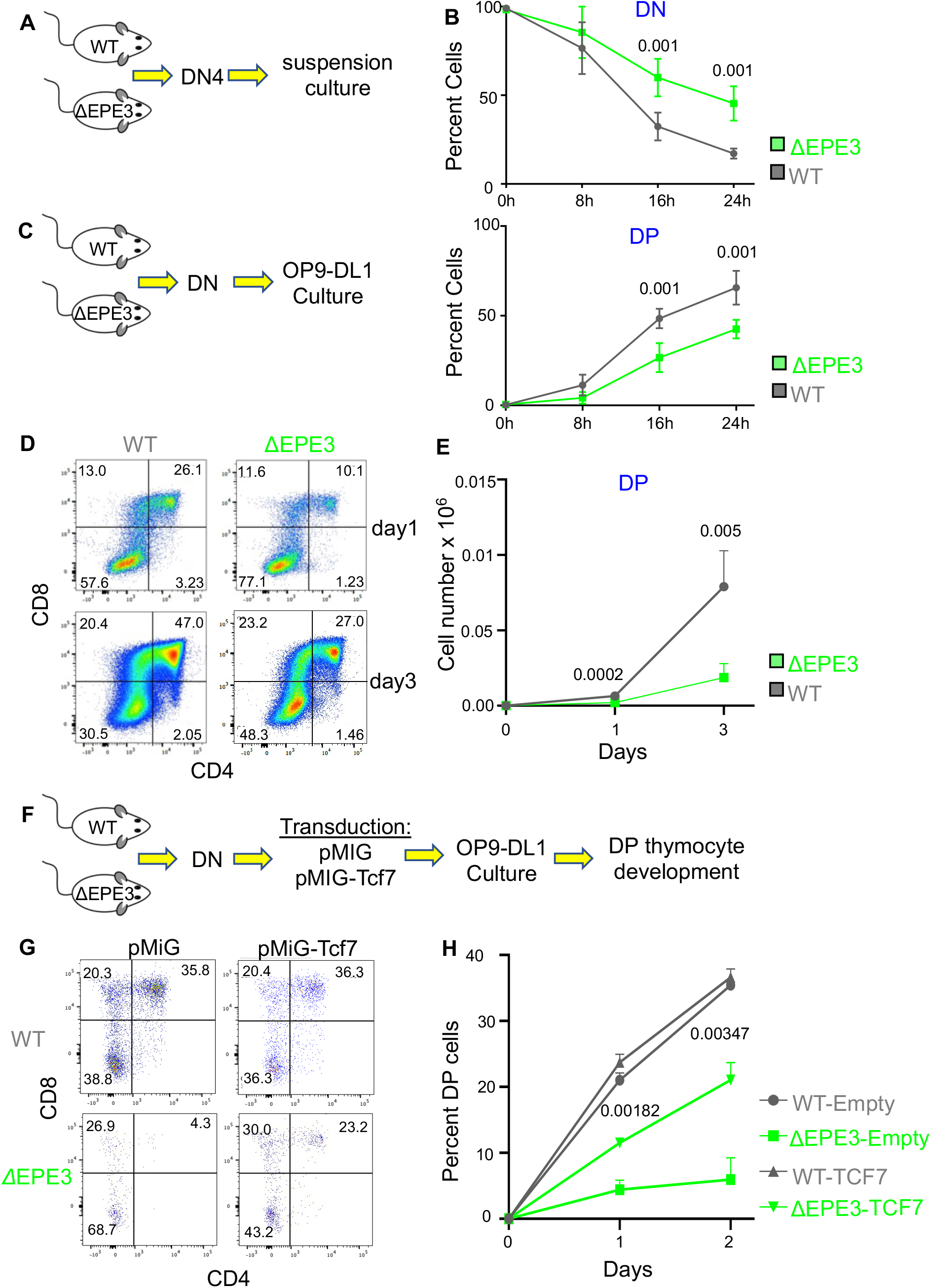
Ablation of EPE3 impairs the development of αβ lineage T cells to the DP stage. **(A)** Schematic representation of the suspension culture experimental design. **(B)** The frequency of DN (top) and DP thymocytes (bottom) was determined by flow cytometry at the indicated times of culture and mean +/- standard deviation (SD) of triplicate measurements was depicted graphically. **(C)** Schematic representation of OP9-DL1 culture experiment. **(D, E)** Development of DN progenitors from WT and ΔEPE3 to the DP stage on OP9-DL1 monolayers was assessed by flow cytometry at the indicated times of culture. Representative histograms of thymic subsets defined by CD4 and CD8 are depicted **(D)** as is a graphical representation (mean +/- SD) of the absolute numbers of DP thymocytes **(E)** on the indicated days of culture. **(F)** Schematic of the TCF1 rescue experiment. **(G)** DN thymocytes isolated as in **(C-E)** were transduced with empty vector (pMiG) or TCF1 (pMiG-*Tcf7)* and cultured in triplicate for 2d on OP9-DL1 monolayers, following which the fraction of DP thymocytes among transduced cells was determined by electronic gating on GFP. Representative two-color histograms of CD4 and CD8 staining are depicted. The mean +/- SD of DP thymocytes for triplicate samples after 1 and 2 days of culture is depicted graphically. Results above are representative of at least 3 experiments performed. p values are indicated.

Because Hi-ChIP analysis suggested that the EPE3 element could interact with other genomic targets, such as the long-distance interaction with the *Skp1* gene (Fig. 1B), we wished to confirm that impaired development of αβ lineage progenitors to the DP stage in ΔEPE3 mice was primarily due to reduced TCF1 expression. For this, we ectopically expressed TCF1 in WT and ΔEPE3 DN thymocytes and assessed their developmental potential on OP9-DL1 monolayers (Fig. 3F). WT DN cells progressed to DP stage over 2 days of culture and were not affected by ectopic expression of TCF1 (Fig. 3G, H). By contrast, ectopic expression of TCF1 significantly rescued the developmental defect of ΔEPE3 DN thymocytes (Fig. 3G, H), demonstrating that the predominant cause of developmental arrest for ΔEPE3 DN thymocytes was reduced expression of TCF1. Collectively, these data show that relatively small changes in the levels of TCF1 expression in thymocytes profoundly affect αβ lineage development.

### EPE3 deletion markedly alters the transcriptomes of ΔEPE3 DN subsets

To gain insight into the mechanisms by which the EPE3 deletion impaired the DN to DP transition, we performed single cell RNA seq (scRNA-Seq) on isolated DN cells from the WT and ΔEPE3 mice (Suppl. Fig. 7). Unsupervised clustering based on differential gene expression resolved DN cells into eight clusters (Fig. 4A, B). All major DN subsets were identifiable in these clusters (Fig. 4A, B) and as expected from the intracellular staining for TCF1 protein, *Tcf7* mRNA was reduced in all the clusters, including the heterogeneous DN1 subset (Suppl. Fig. 8A). Consistent with flow cytometric analysis, the proportion of DN3 cells increased in ΔEPE3 mice. Among DN3 cells, we observed two distinct sub-populations in WT and ΔEPE3 thymocytes, referred to as WT-bias and ΔEPE3-bias, respectively (Fig. 4A, B). We observed that proportions of cells in the DN2, DN3-ΔEPE3-biased and DN4 clusters were increased in ΔEPE3 mice compared to WT (Fig. 4C). Together, these data indicate that reduced TCF1 expression caused by EPE3 deletion begins to alter the transcriptomes of thymic progenitors beginning in DN2 cells and continuing to the DN4 stage.

**Figure 4:**
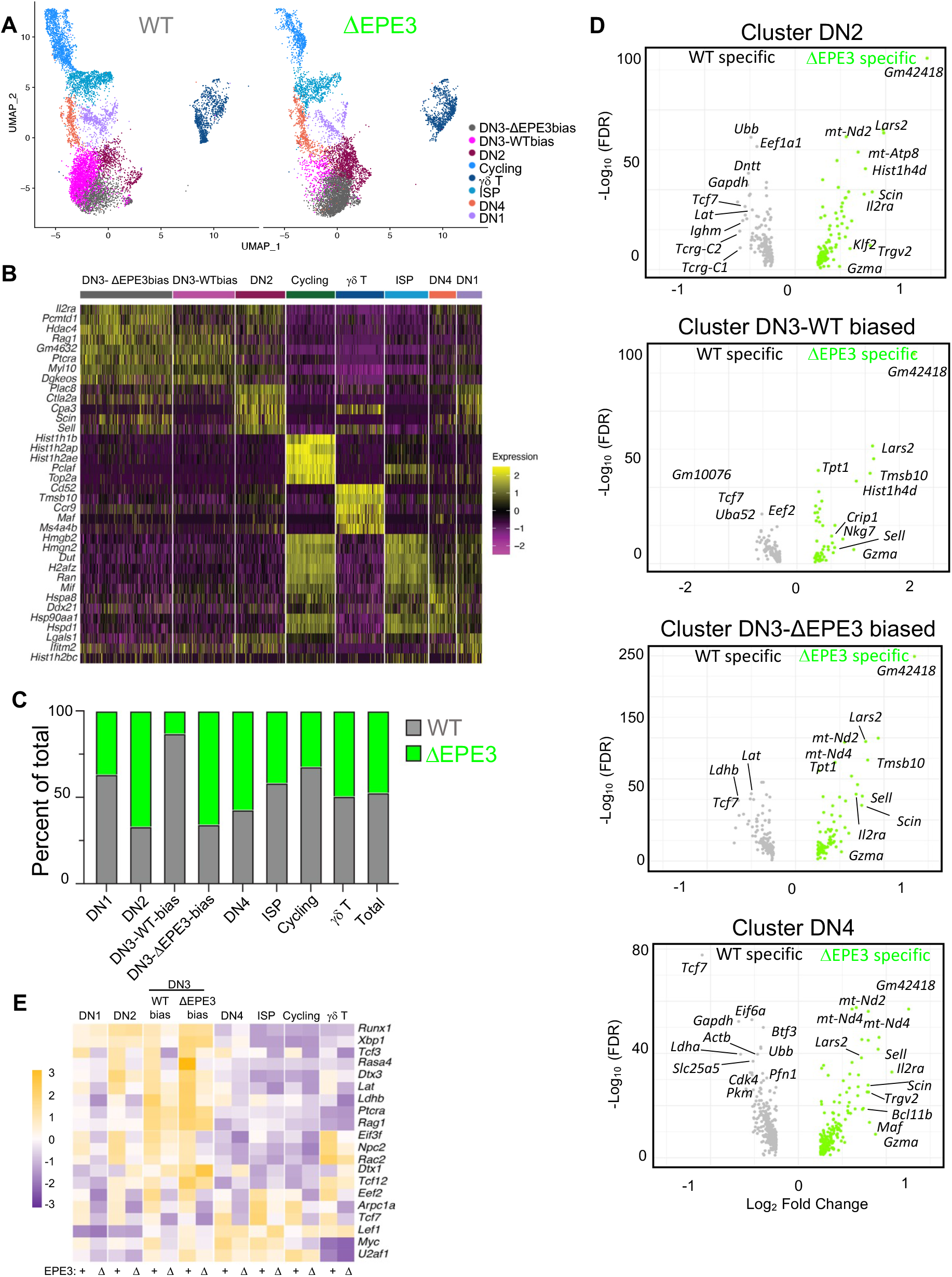
ScRNA-Seq analysis of DN thymocytes from ΔEPE3 mice. **(A)** UMAPs of clusters identified by scRNA-Seq analysis of DN thymocytes isolated from WT and ΔEPE3 mice. **(B)** Heat map representation of genes used to define the clusters. **(C)** Bar graph illustrating relative proportions of each cluster among DN thymocytes from WT and ΔEPE3 mice. **(D)** Volcano plots displaying differentially expressed genes in DN2, DN3-WT biased, DN3-ΔEPE3 biased, and DN4 clusters. **(E)** Heat map representation of expression changes of selected genes, including TCF1 targets, in all clusters.

To investigate the basis for the developmental impairment observed in the ΔEPE3 mice, we identified differentially expressed genes (DEGs) in the DN2, DN3, and DN4 clusters (Fig. 4D, E). Amongst DEGs, we observed previously identified TCF1 targets including *Lef1, Dtx1, Ptcra,* and *Lat* ^10^ (Fig. 4D, E and Suppl. Fig. 8B). In addition, there were DEGs that had not been previously implicated as TCF1 targets such as *Arpc1a, Ldhb, Myc,* and *Tcf3* (E2A) (Fig. 4E and Suppl. Fig. 8B). TCF1 ChIP-Seq data from DP thymocytes revealed TCF1 binding near all of these genes, raising the possibility that these genes are also direct TCF1 targets ^2^. Interestingly, a number of these putative TCF1 target genes encode regulators of metabolism including *Ldha, Ldhb*, and *Gapdh*, which were all downregulated among DN2-4 cells from ΔEPE3 mice (Fig. 4D, E and Suppl. Fig. 8B). Together, these data reveal that the DN2 to DN3 transition and the capacity of the pre-TCR to promote the DN to DP transition is likely to be compromised in ΔEPE3 mice by reduced expression of essential molecular effectors of signaling and gene regulation (*Ptcra, Lat, Myc,* and *Lef1)*, induction of inhibitors of Notch signaling (Dtx1) (Fig. 4D, E), and metabolic changes associated with cell proliferation ^5, 32, 33, 34^.

### Notch binding site (NBS) and EPEs play distinct roles in regulating TCF1 expression

Pseudobulk ATAC seq analysis on WT and ΔEPE3 T cell progenitors showed that all 5 EPEs were accessible in DN T cell progenitors (Fig. 5A). Previous analysis of regulatory elements controlling the *Tcf7* locus implicated a 1kb region of chromatin that encompassed the 207bp of EPE3 but extended further to include a NBS located 177bp away from the EPE3 deletion (Fig. 5A and boxed region in Suppl. Fig. 9) ^10, 21^, raising the possibility that both E-protein binding to EPEs and Notch binding to the NBS were required for *Tcf7* expression. Because the EPE3 deletion preserved the NBS (Suppl. Fig. 1A), we sought to determine the contribution of Notch function in regulating TCF1 levels in ΔEPE3 progenitors. First, ChIP analysis using anti-Notch1 showed that the EPE3 deletion did not disrupt Notch binding to the NBS near EPE3 (Fig. 5B). Second, we determined if ectopic expression of the active intracellular fragment of Notch1 (ICN1) ^35, 36^ could regulate TCF1 expression levels and whether this was affected by the EPE3 deletion (Fig. 5C). To this end, we retrovirally transduced fetal liver DN precursors from WT and ΔEPE3 mice with ICN1 and evaluated the effect on TCF1 expression by intracellular staining (Fig. 5C-E). ICN transduction of CD25^-^ DN1 progenitors induced TCF1 expression among TCF1^-^ progenitors, eliminating the TCF1^-^ subpopulation in both WT and ΔEPE3 cells, thereby demonstrating that the capacity of ICN1 to induce TCF1 expression was not impaired by the ΔEPE3 allele (Fig. 5D-F). Importantly, the TCF1 expression levels among cells that expressed TCF1 prior to ICN transduction (TCF1^+^) were reduced in progenitors from ΔEPE3 mice expressed (Fig. 5D, E, G; MFI plots). Moreover, ICN1 transduction failed to restore TCF1 expression by ΔEPE3 progenitors to the level observed in WT progenitors (Fig. 5D, E, G; MFI plots). Together, these data show that the EPE3 element is dispensable for Notch-mediated initiation of TCF1 expression but regulates the levels of TCF1 expression thereafter through E protein activity.

**Figure 5:**
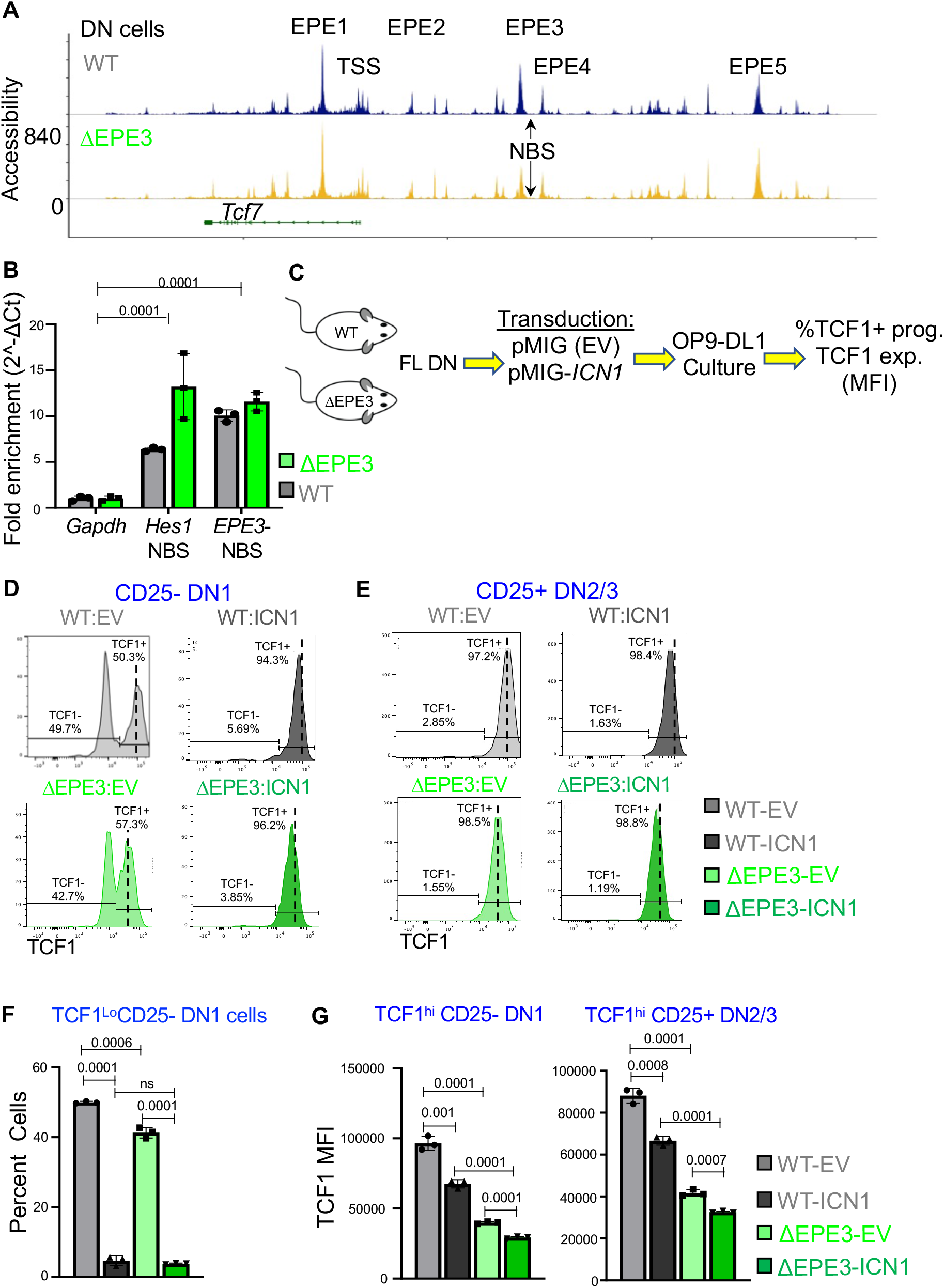
Role of Notch in regulating TCF1 expression. **(A)** ATAC-seq analysis of chromatin accessibility near the *Tcf7* locus in DN thymocytes from WT and ΔEPE3 mice. A 90kb interval of Chr11 is depicted. **(B)** ChIP qPCR analysis of Notch binding to the *Gapdh, Hes1 NBS,* and *EPE3-NBS* elements. *Hes1* serves as a positive control for Notch binding and *Gapdh* served as a negative control. The ChIP signals for *Hes1* and *EPE3-NBS* were normalized to *Gapdh*. **(C)** Schematic for the Notch rescue experiment. **(D, E)** The ability of activated Notch to modulate TCF1 expression was assessed by retroviral transduction of WT or ΔEPE3 fetal liver precursors with empty vector (EV; pMiG) or the active intracellular fragment of Notch1 (pMiG-*ICN1*) after 4 days of culture on OP9-DL1 monolayers. After 3 additional days of culture the effect of activated Notch on TCF1 expression was assessed by intracellular flow cytometry on electronically gated GFP+ CD25- DN1 progenitors **(D)** or CD25+ DN2/3 progenitors **(E)**. Representative histograms are depicted. Dashed lines marks the mean peak intensity for the TCF1 in WT control transduced TCF1^+^ cells. ICN transduction reduces the intensity of intracellular TCF1 staining. **(F)** The fraction of TCF1low/negative cells in triplicate technical replicate cultures of the CD25- DN1 progenitors above was quantified and depicted graphically as the mean +/- SD. **(G)** The level of TCF1 expression was assessed by determining the MFI of TCF1 expression by flow cytometry of the TCF1hi fraction of triplicate cultures of both CD25- DN1 and CD25+ DN2/3 progenitors. Results are representative of 3 experiments performed. p values are shown.

## Discussion

Because of the critical role of TCF1 in all stages of T cell development and peripheral differentiation, the mechanisms controlling its expression are of significant interest. Previous efforts to understand the *trans* acting factors and *cis* elements controlling the *Tcf7* locus focused on the role of Notch signaling, given its ability to induce TCF1 expression during T lineage commitment. Here we provide new insights into the Notch-independent role of the E protein axis and the requisite genomic elements through which this axis controls TCF1 expression.

A recurring theme in hematopoiesis is activation of lineage-specific gene expression and repression of lineage-inappropriate gene expression to commit multipotential progenitor cells to the differentiation program of a specific cell type. TCF1 and its targets *Bcl11b* and *Gata3*, serve as critical molecular effectors of commitment of thymic progenitors to the T lineage ^10, 37^. *Tcf7*, the gene that encodes TCF1, is activated in response to Notch signals provided by the thymic microenvironment ^37^. We provide evidence that the E box DNA binding protein family members expressed by thymocytes (E2A and HEB), play an essential role in maintaining and determining the optimal level of TCF1 expression starting at DN2 stage. There are five E protein bound elements (EPE) near the *Tcf7* locus that play distinct roles in supporting TCF1 expression in the γδ and αβ T lineages. EPE1, and 5 selectively regulate TCF1 expression only in γδ T cell progenitors, while EPE3 controls TCF1 levels in both αβ and γδ T cells. Because ablation of EPE3 eliminates the E-protein binding sites, but retains adjacent Notch binding sites, the *Tcf7* locus remains Notch-responsive. Consequently, we hypothesize that Notch signaling is responsible for initiating TCF1 expression among early T cell progenitors (ETP), but then maintenance of TCF1 expression and determination of expression levels is transferred from Notch to E-proteins upon arrival at the DN2 stage and beyond.

A particularly interesting aspect of our analysis is the finding that a reduction in TCF1 expression of less than two-fold in ΔEPE3 thymocyte causes a severe impairment in αβ T cell development, further emphasizing that the level of TCF1 expression plays a critical role in controlling developmental transitions. This may also indicate why ectopic expression of TCF1 fails to fully rescue T cell development, as maintaining the precise TCF1 dosage is challenging through ectopic methods. Because TCF1 regulates numerous target genes through its ability to act as a pioneering factor ^2^, the developmental impairment likely results from dysregulation of numerous genes. Among these are several that are noteworthy, including reduced expression in ΔEPE3 progenitors of *Lat* and *Ptcra*, which should impair pre-TCR signaling ^10, 38^, and induction of *Dtx1,* which has been reported to interfere with the ability of Notch signals to support development ^34^. Moreover, key DNA binding proteins that support the pre-TCR induced DN to DP transition are also reduced in ΔEPE3 progenitors, including *Myc* and *Tcf3* (E2A) ^2, 33^. Finally, several metabolic regulators (*Gapdh*, *Ldha* and *Ldhb*) were down-regulated in ΔEPE3-DN cells. Because maintenance of glycolysis by Notch is important for development to the DP stage ^39^, and because the accumulation of lactate blocks T cell proliferation ^40^, these changes in gene expression may also contribute to metabolic alterations that attenuate development. While these changes in gene expression provide potential explanations for the attenuation of αβ T cell development, the targets through which the combined reduction of TCF1 and E protein activity facilitate γδ lineage commitment remain undefined.

EPE3 can be characterized as a classic enhancer that augments TCF1 expression by associating with the *Tcf7* promoter and transcription start site. Several transcription factors have been shown to bind to this region in addition to E proteins and Notch. The ΔEPE3 deletion described here removes putative E protein binding sites but leaves intact the Notch binding site. The profound effect of ΔEPE3 on αβ T cell development clearly demonstrates the importance of E protein binding to EPE3 and the insufficiency of Notch binding to the adjacent NBS to maintain or restore optimal TCF1 expression. Indeed, even ectopic expression of ICN fails to retore TCF1 expression in ΔEPE3 DN cells to levels seen in wild type progenitors, suggesting that the critical role of E-proteins is manifest on a fully Notch-responsive *Tcf7* allele. Unlike the effects of deleting E-protein binding sites seen here, previous studies show that deleting NBS adjacent to EPE3 does not significantly affect αβ T cell development^21^. We surmise that absence of that NBS can be compensated by Notch binding to other presently uncharacterized regulatory sequences.

These observations raise the question as to how residual TCF1 expression is maintained from ΔEPE3 alleles. One possibility is that *Tcf7* transcription is regulated by sequences flanking the E-protein binding sites in the EPE3 region. In addition to an NBS, the flanking region also has predicted binding sites for GATA3, TCF1 and Runx. A related study that ablated a larger genomic interval encompassing the EPE3 as well as the TCF1, Gata3, and Runx sites, observed greater reduction in TCF1 expression than did ablation of EPE3 alone, suggesting that these flanking sequences may be responsible for supporting residual expression of TCF1 in EPE3 mutant cells^21^. In support of this hypothesis, we found that chromatin accessibility of this region was reduced, but not eliminated, on ΔEPE3 alleles. Alternatively, it is possible that another EPE, such as EPE5, sustain the lower levels of *Tcf7* expression observed in ΔEPE3 progenitors. Although individual ablation of the other EPEs did not majorly impact αβ T cell development, it remains possible that they may do so when combined with EPE3. EPE5 is of particular interest, because we observed contacts between EPE5 and the *Tcf7*-TSS in Hi-ChIP analysis. These contacts were weaker than the EPE3-TSS contacts, but EPE5-TSS contacts were further reduced in ΔEPE3 progenitors, raising the possibility of cooperation.

During T cell development, E-protein function is dynamically regulated post-transcriptionally through the induction of members of the antagonistic Inhibitor of DNA binding (Id) family ^23, 24, 41,42^. Id3 is induced in proportion to TCR signaling intensity and duration and blocks E-protein function ^23, 24, 43^. Accordingly, the activity of E-proteins and consequently TCF1 expression are responsive to the nature of TCR signals transduced, with more intense signals leading to greater repression of TCF1 expression ^16^. In support, we recently demonstrated that the weak TCR signals that lead to αβ lineage commitment preserves TCF1 expression levels, while the more intense TCR signals that promote γδ lineage commitment result in reduced TCF1 expression, which acts to facilitate γδ lineage commitment ^16^. Nevertheless, because Id3 is regulated by the Egr family of transcription factors, which are induced by a diverse array of stimuli, E-protein activity is almost certainly also controlled by extracellular signals in addition to those emanating from the TCR. Finally, our finding that E protein binding to EPE3 plays a key role in regulating TCF1 expression in developing thymocytes has implications beyond T cell development. For example, TCR modulation of E-protein binding to EPE3 may regulate the expression of TCF1 in peripheral T cells, where it has been shown to play a critical role in effector function in the context of exhaustion due to prolonged TCR stimulation, such as is observed in chronic viral infection or in cancer ^44, 45, 46, 47^. Indeed, the ability to express TCF1 by “stem-like” precursor exhausted (Tpex) T cells, which can be reinvigorated by checkpoint blockade in cancer, may be controlled through by the TCR-Id3-E-protein axis and EPE3 ^46, 47^.

## MATERIALS AND METHODS

### Mice, tissue and cells extraction

Wild type male and female C57BL/6J mice were obtained from the Jackson Laboratory (Bar Harbor, ME). ΔEPE1, ΔEPE2, ΔEPE3, ΔEPE4 and ΔEPE5 mice were generated using CRISPR-Cas9 technology as described in ^48^. gRNAs sequences and primers for genotyping are listed in Tables 1 and 2. ΔEPE1 mice were generated as described^16^. Generation of *Tcf7*-/- mice is previously described^5^. Mice were bred and maintained either in the animal facility at the National Institute on Aging (NIA) or at Fox Chase Cancer Center (FCCC). All the mice used in the experiment were age-matched (8-14 weeks old). The studies were carried out in accordance with the recommendations in the Guide for the Care and Use of Laboratory Animals. Thymi were isolated and gently pressed through 70 *µ*M strainer to extract the thymocytes. The thymocytes were resuspended in 10 ml of RPMI medium with 2% FBS (Gibco, MD).

**Table-1.**
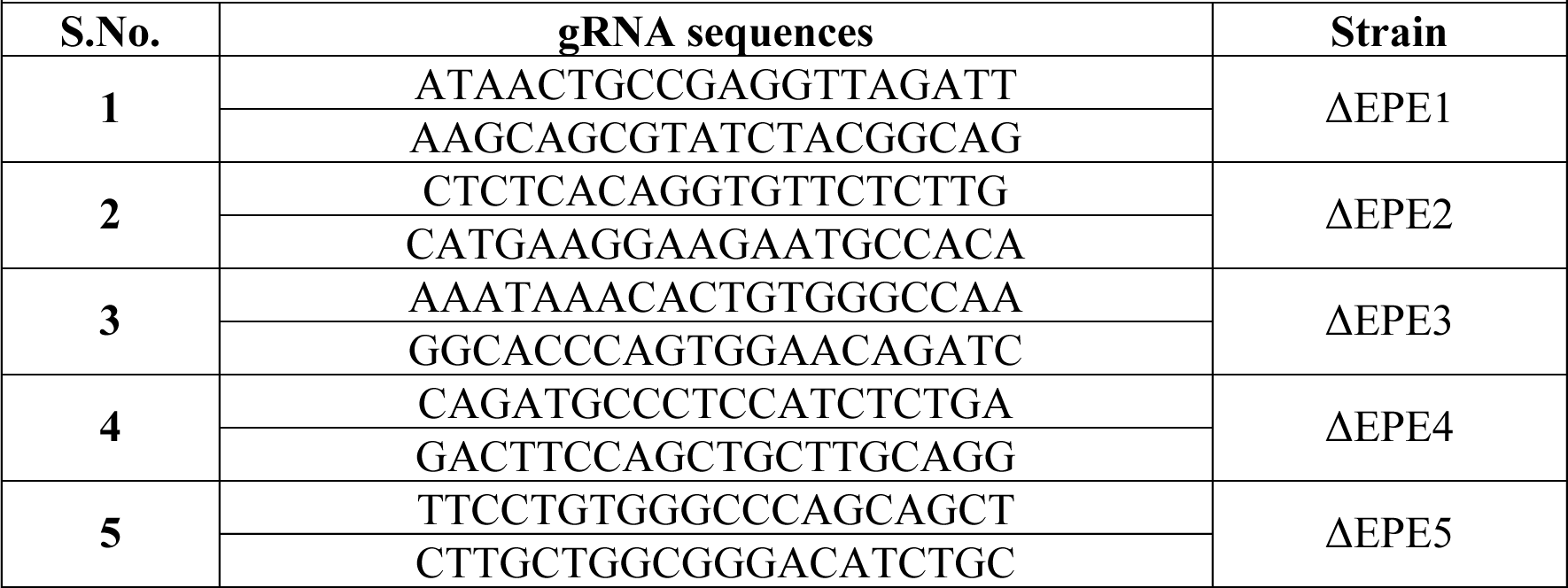
List of gRNA sequence for CRISPR mice.

**Table-2.**
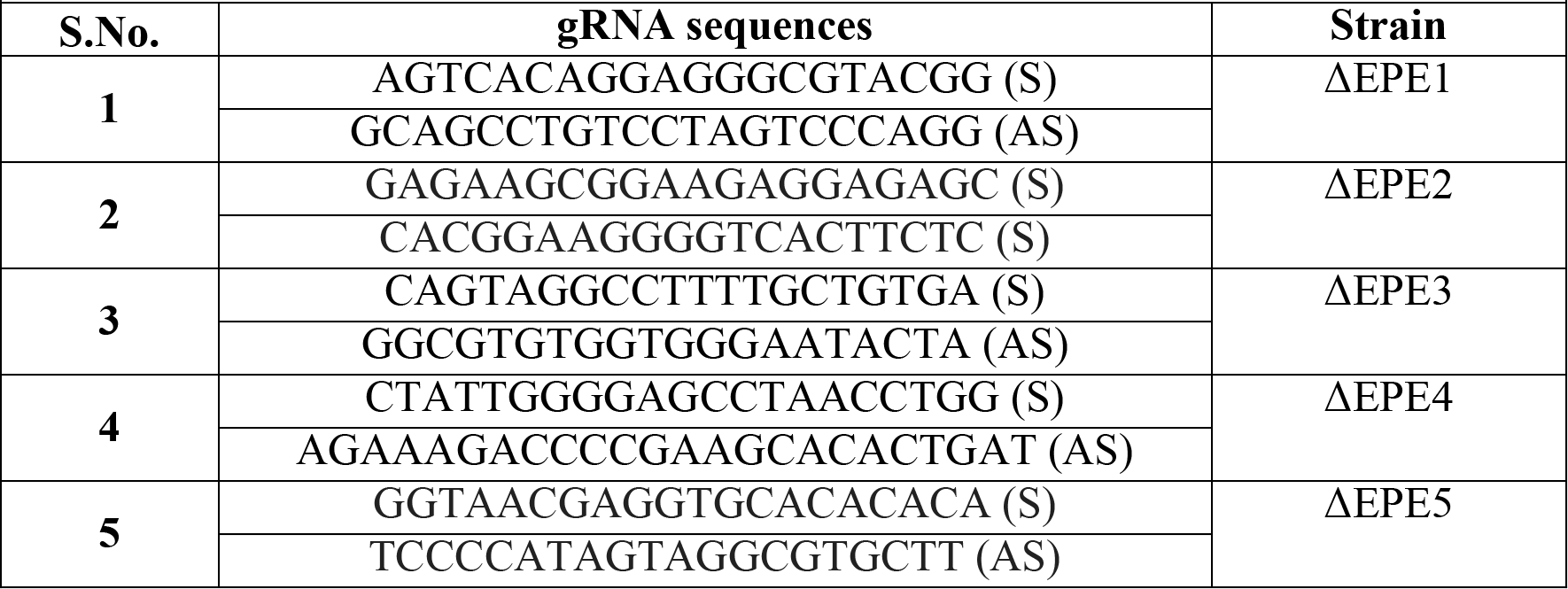
List of genotype primers sequence for CRISPR mice.

### Flow cytometry and Cell Sorting

Thymocytes were stained and acquired on Symphony-analyzer (Becton-Dickinson, DC.) and analyzed with FlowJo version 10.0 (Flowjo LLC, USA). eBioscience^TM^ Fixable Viability Dye eFluor-780 (ThermoFisher Scientific, MA) was used to exclude dead cells. Fluorescent antibodies were purchased from from BD Biosciences, eBioscience or BioLegend and listed in Table-3 and other reagents used for FACS are listed in supplementary (Table-4). In brief, cells were incubated with FC block and stained with antibodies for surface marker and fixed with 2% paraformaldehyde. For intracellular staining, cells were permeabilized using the eBioscience^TM^ FoxP3/transcription factor staining buffer set (ThermoFischer Scientific, MA) and stained with antibodies against the transcription factors. For sorting, CD8α cells were removed by incubating thymocytes with CD8α microbeads (Miltenyi Biotech, USA) as per user’s manual. CD8-depleted cells were stained with antibodies for surface markers and resuspended in RPMI-1640 supplemented with 2% FBS. Cells were sorted in RPMI-1640 medium supplemented with 10% FBS.

**Table-3.**
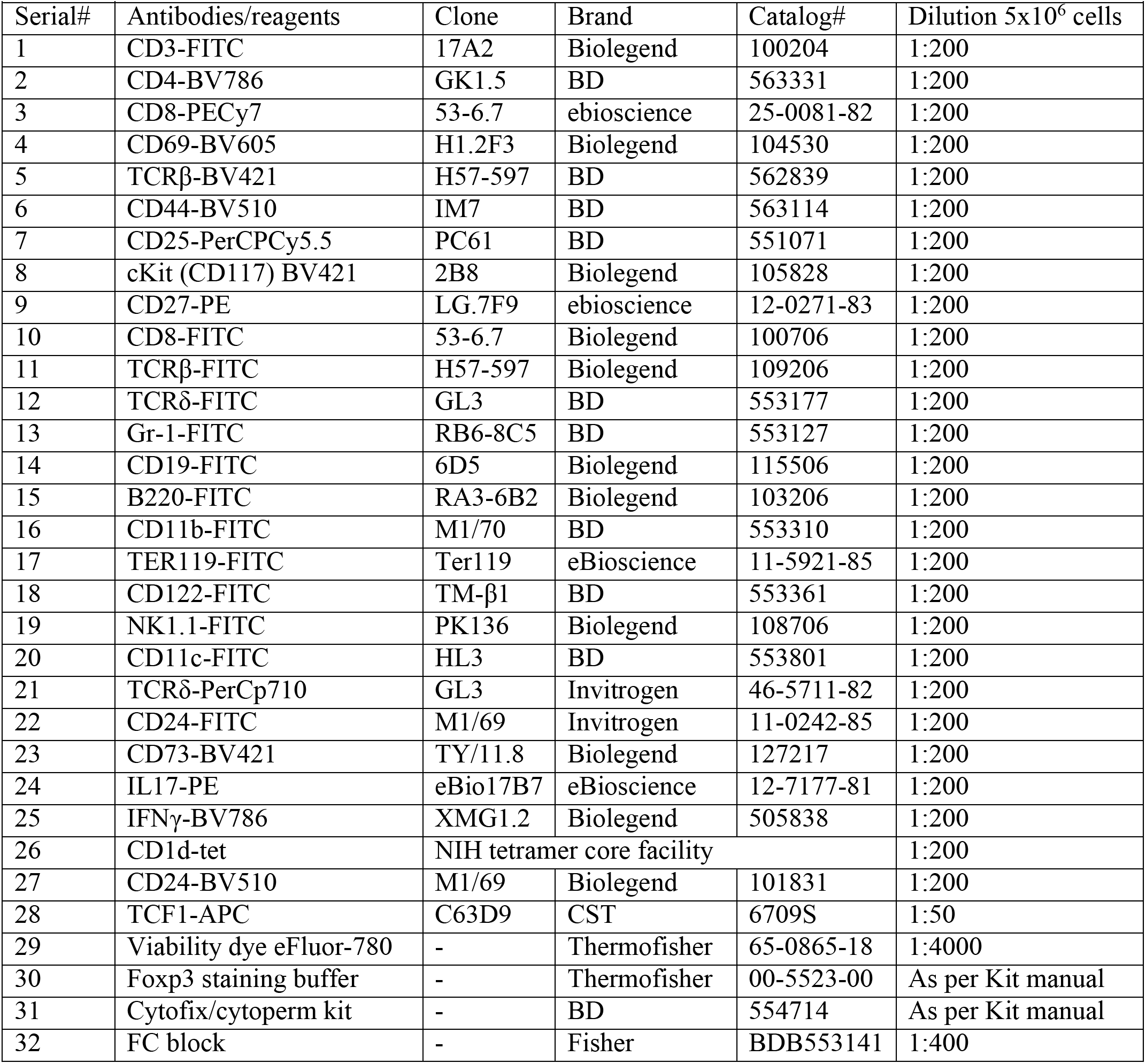
List of antibodies against surface markers or intracellular proteins/FACS reagents.

**Table-4.**
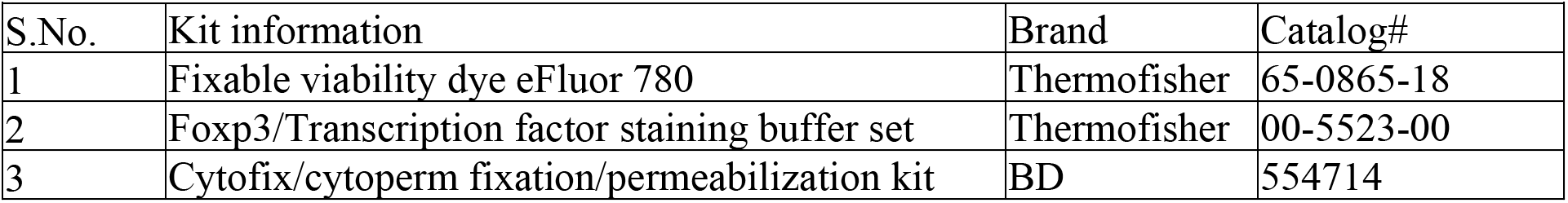
List of kits used in FACS.

### In vitro culture of DN progenitors

CD8-depleted cells were stained with antibodies for surface markers. Cells were resuspended and sorted in RPMI-1640 medium supplemented with 10% FBS. DN4 thymocytes (Lineage-depleted with antibodies reactive to: TCRβ, CD4, and CD8, CD11c, CD11b, Gr1, CD19, B220, Ter119) were isolated by flow cytometry based on the absence of expression of CD44 and CD25, and cultured in 12-w plates overnight at 10^6^ cells/ml at 37֯C with 5% CO2. Double positive (CD4+CD8+) thymocytes were quantified by flow cytometry every 8h up to 24 hours. For culture of adult DN thymocytes on OP9-DL1 cultures, DN thymocytes were isolated by magnetic bead depletion using anti-CD4 and anti-CD8 magnetic beads, following which they were plated on OP9- DL1 monolayers as 100,000 cells per well of a 24 well plate in the presence of IL-7 and Flt3 as described ^49^. After 1 and 3days of culture, development to the DP stage was assessed by flow cytometry. For retroviral transduction, the DN thymocytes were spin infected with retrovirus encoding empty vector (pMiG) or *Tcf7* (pMiG-*Tcf7)* in the presence of 8μg/ml polybrene for 90min, following which the behavior of transduced progenitors was monitored by flow cytometry by electronic gating on GFP+ progenitors at days 1 and 2 post-tranduction ^16^. For transduction of fetal liver precursors, embryonic day 14 (E14) fetal liver precursors from WT or ΔEPE3 mice were cultured on OP9-DL1 monolayers for 4d as described ^16^, transduced with empty vector (pMiG) or the Notch intracellular domain (pMiG-*ICN1)*, cultured for 3 days and then subjected to intracellular staining with anti-TCF1 antibody as described above.

### ChIP and HiChIP

E14 fetal liver precursors from WT and ΔEPE3 were cultured on OP9-DL1 monolayers for 7 days, following which DN3 cells were isolated by flow cytometry, fixed in 1% formaldehyde for 10 minutes, and sonicated to produce chromatin fragments, as described ^16^. The resulting fragments were immunoprecipitated with 5 μg of anti-Notch antibody (Cell Signaling Technology #3608) with Protein G Dynabeads (Invitrogen 10003D). Eluted chromatin fragments were analyzed in triplicate by quantitative PCR (qPCR) using SYBR Green and primer sets spanning sites in the *Gapdh, Hes1,* and *EPE3* (*Gapdh – F,* TGGCGTAGCAATCTCCTTTT*; R,* TGGCGTAGCAATCTCCTTTT*; Hes1 – F,* CGTGTCTCTTCCTCCCATTG*; R,* CGTGTCTCTTCCTCCCATTG*; EPE3 – F,* CACCGAGCATTCTCAGCAGCA*; R,* ACAGCTTTTATCGCACGTTTATGAAGG). qPCR results from the ChIP analysis were normalized to *Gapdh* and represented graphically as the mean +/- SD of triplicate measurements. To prepare cells for HiChIP, DN3 precursors were isolated by flow cytometry following culture on OP9-DL1 monolayers for 7 days as described above. HiChIP assays were conducted using duplicate samples of wild-type pro-B cells, wild-type DN3 cells, and ΔEPE3 DN3 cells. The Arima HiC+ kit (Arima Genomics, Cat. # A101020) was employed, strictly following the manufacturer’s protocol as outlined in Arima-HiC documents A160168 v00 (HiChIP) and A160169 v00 (library preparation using a Swift Biosciences Accel-NGS 2S Plus DNA Library Kit). For each HiChIP sample, 5×10^6^ cells were collected, and a total of 2.5 μg of H3K27ac antibody (Active Motif, Cat. #39133) was utilized. The resulting barcoded HiChIP libraries were combined and subjected to sequencing using an Illumina NovaSeq instrument to produce an average depth of 200 million reads. HiChIP reads were processed using Juicer to generate .hic files^50^. Juicer was run with the flags “-g mm10 -s Arima”. To compare multiple experiments, Hi-ChIP data were normalized by down-sampling to the same number of total reads. Contacts within 2,000 bp and reads with mapping quality scores below 30 were removed prior to down-sampling. To quantify the interaction frequency between specific regions, such as EPE sites and promoters, the number of contacts was counted after the experiments had been down-sampled to the same number of total reads. Hi-ChIP contact maps were visualized using cooltools^2^ and are shown at 1,000 bp resolution. Difference maps were created by subtracting the wild-type contacts from the mutant contacts.

### Single cell RNA-Seq

#### Cell isolation

CD4-CD8-double negative (DN) cells were obtained from C57BL/6 (WT-DN) and ΔEPE3 (ΔEPE3-DN) mice. In brief, thymocytes were depleted using CD8α microbeads (Miltenyi Biotech, USA) (Suppl. Fig. 7A). The CD8-depleted fraction was incubated with lineage marking antibodies (TCRβ, CD4, and CD8, CD11c, CD11b, Gr1, CD19, B220, Ter119) and then lineage-cells were sorted in RPMI-1640 medium supplemented with 10% FBS. Post-sorting, a small fraction of sorted cells was used to check the purity (>98%). 200,000 cells per sample were provided for the library preparation.

#### Library Generation

Using a Chromium Next GEM Single Cell 3ʹ _GEM, Library & Gel Bead Kit v3.1 (10x Genomics, PN-1000121) and the Chromium Chip G Single Cell Kit (10x Genomics, PN-1000120), the thymocytes, at 1,000 cells per *µ*l, were loaded onto a chromium single-cell controller (10x Genomics) to generate single-cell gel beads-in-emulsion (GEMs) according to the manufacturer’s protocol. Briefly, approximately 10,000 cells per sample were added to chip G to create GEMs. Cells were lysed, and the bead captured poly(A) RNA was barcoded during reverse transcription in Thermo Fisher Scientific Veriti 96-well thermal cycler at 53 °C for 45 min, followed by 85 °C for 5 min. cDNA was generated and amplified. Quality control and quantification of the cDNA were conducted using Agilent’s High Sensitivity DNA Kit (5067- 4626) in the 2100 Bioanalyzer. scRNA-seq libraries were constructed using a Chromium Single Cell 3ʹ Library Kit v3.1 and indexed with Single Index Kit T Set A (10x Genomics, PN-1000213). The libraries were sequenced using an Illumina NovaSeq 6000 sequencer with a paired-end, single-indexing strategy consisting of 28 cycles for read 1 and 91 cycles for read 2.

#### Data processing

Demultiplexing of raw base call files into FASTQ files was completed with 10X Genomics Cell Ranger (v.3.0.2) mkfastq coupled with mouse reference version mm10. The results from Cell Ranger were processed using DoubletFinder and Seurat, following standard procedures ^51, 52, 53^. DoubletFinder was used on each sample individually to remove doublets. One WT and one PK3 mutant sample were paired together and integrated with a second pair of WT and PK3 mutant samples to remove experimental batch effect. Integrated data were clustered with a resolution of 0.4. Clusters comprising less than 5% were removed from further analysis, as were NKT cells. Differentially expressed genes between genotypes were identified with the *FindMarkers* function of Seurat.

### Single-cell ATACseq Analysis

#### Cell isolation

Sorted WT-DN and ΔEPE3-DN cells isolated as above were used for single cell ATAC seq analyses. For each sample, 100,000 cells were used for nuclei isolation as described^54^. Briefly, 100,000 cells were centrifuged at 500 g for 5 minutes at 4°C. Cell pellet was washed with 50 μl ice-cold PBS. Cell pellet was gently resuspended in 50 μl cold lysis buffer (10mM Tris-Cl, pH 7.4, 10 mM NaCl, 3mM MgCl2, 0.1% IGAPAL CA-630) and centrifuged at 500 g for 10 minutes at 4°C. Isolated nuclei were passed through Flowmi Cell Strainer (40 μM) (Scienceware, H13680-0040).

#### Library Generation

Using the Chromium Next GEM Single Cell ATAC Library & Gel Bead Kit v1.1 (10x Genomics, PN-1000175) and the Chromium Next GEM Chip H Single Cell Kit v1.1 (10x Genomics, PN-1000161), thymocyte nuclei, at 7,000 nuclei per *µ*l, were loaded onto a Chromium controller (10x Genomics) to generate single-nuclei gel beads-in-emulsion (GEMs) after which library construction is performed according to the manufacturer’s protocol. Briefly, nuclear DNA is fragmented at open regions of the chromatin with transposase while adapter sequences are added at the ends. The fragmented nuclear DNA is then loaded on H chips and partitioned into nanoliter-scale GEMS (Gel Beads-in-emulsion). Illumina P5 sequence, a barcode and read 1 sequence are added by PCR amplification. Unused primers and reagents are removed from the sample by SPRI beads. The library construction is completed by the addition of both P7 and a sample index. The libraries were sequenced using an Illumina NovaSeq 6000 sequencer with a paired-end, dual-index strategy consisting of 50 cycles for read 1, 50 cycles for read 2, 8 cycles for the i7 index and 16 for the i5.

#### Data processing

Demultiplexing of raw base call files into FASTQ files was completed with 10X Genomics Cell Ranger-ATAC MKFASTQ coupled with mouse reference version mm10. The results from Cell Ranger were processed using Seurat V3 and Signac V0.2.1 and 1.0.0, following standard procedures ^53, 55^. Genomic traces were generated from single-cell ATAC data that has been converted to pseudo-bulk tracks of DN cells, as described^55^.

### Software and Statistics

Statistical significance was determined by the student’s unpaired *t*-test.

## AUTHOR CONTRIBUTIONS

JMS and DLW conceived and designed the study. JMS, DW and RS oversaw the study and wrote the manuscript. AV, BA, FM, CAS, SS and AC performed experiments and analyzed data. XQ, BK contributed to the generation and characterization of EPE mice. LC, C M-M, SD, NO, RB and RS contributed critical expertise.

## COMPETING FINANCIAL INTERESTS

The authors have no competing interests.

## ACKNOWLEDGEMENTS

We gratefully acknowledge the assistance of the core facilities of FCCC: Cell Culture, DNA Sequencing, Flow Cytometry, and Laboratory Animal. DLW was supported by NIH grant P01AI102853, core grant P30CA006927, the Bishop Fund and an appropriation from the Commonwealth of Pennsylvania. We acknowledge help from Dr. Fatima Braikia and NIA-CMS. This work was supported in part by the Intramural Research Program of the National Institute on Aging.

---

**Supplementary Figure 1:**
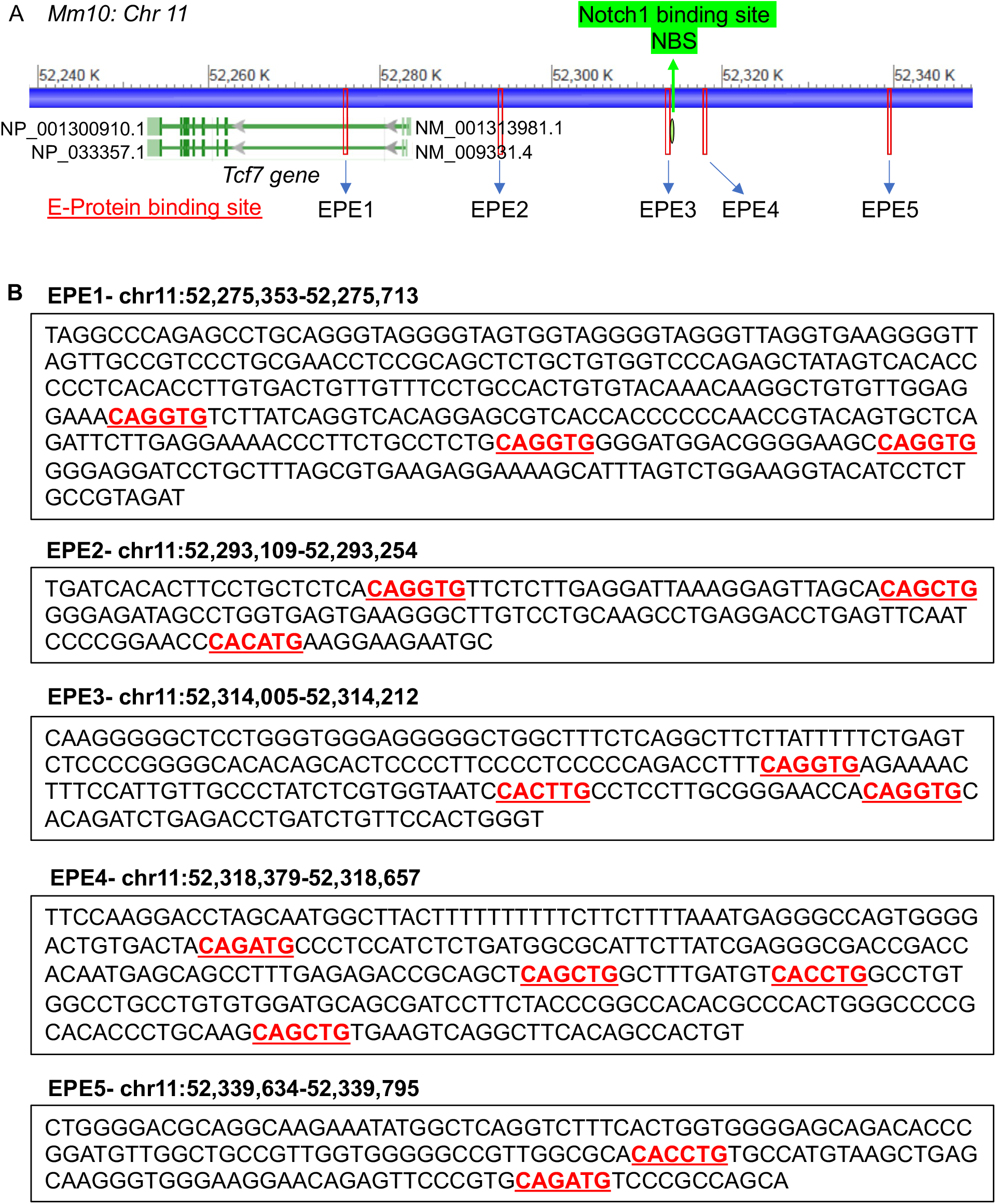
CRISPR-deletion of E-protein binding elements (EPE) 1, 2, 3, 4, and 5. **(A)** Schematic depicts the location *Tcf7* gene, ChIP-seq peaks defining EPE1, EPE2, EPE3, EPE4, and EPE5 clusters, and the Notch binding site (NBS) adjacent to EPE3 (Mm10). **(B)** Sequences deleted in EPE1, EPE2, EPE3, EPE4, and EPE5 using CRISPR-Cas9 system. Consensus E protein binding sites (CANNTG) are bolded and underlined in red.

**Supplementary Figure 2:**
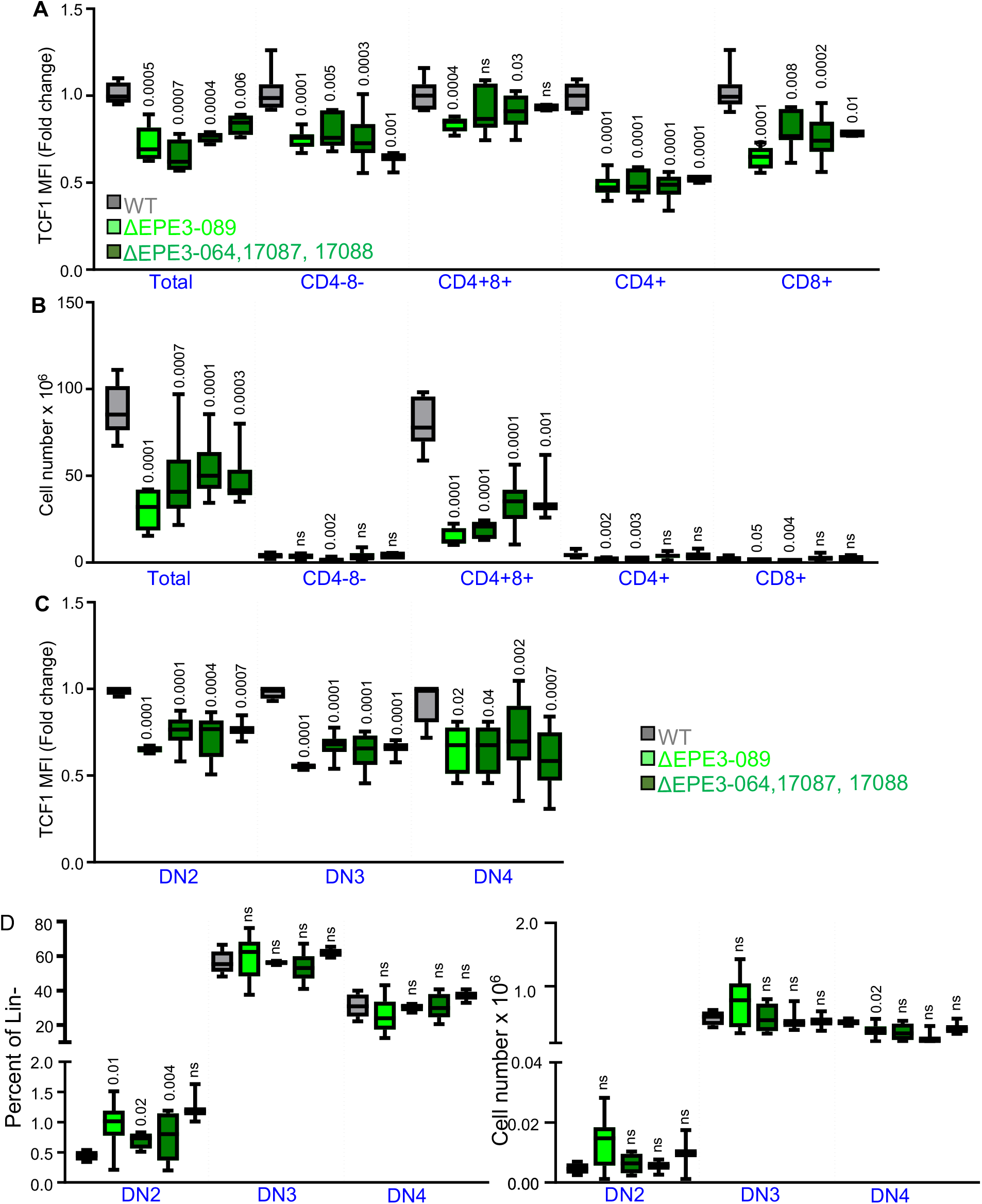
Impact of ablation of EPE3 on development of αβ lineage thymocytes in other ΔEPE3 founder lines. **(A)** Graphical representation of the fold-change in TCF1 expression as determined by intracellular staining. The following founder lines were analyzed: WT (n=10), ΔEPE3-089 (n=7), ΔEPE3-064 (n=8), ΔEPE3-17087 (n=10), and ΔEPE3-17088 lines (n=6). Cells from ΔEPE3-089 line were used for majority of the assays presented in this study. **(B)** The absolute number of thymocytes in the indicated thymic subsets is depicted graphically in the same mice as in **(A)**. **(C)** Graphical representation of the fold-change in TCF1 expression as determined by intracellular staining of the DN subsets from WT and the ΔEPE3 founder lines in **(A)**. **(D)** Frequency (left) and absolute number (right) of the indicated DN subsets from WT and the ΔEPE3 founder lines in **(A).** Results are representative of 3 or more experiments performed. p values are indicated.

**Supplementary figure 3:**
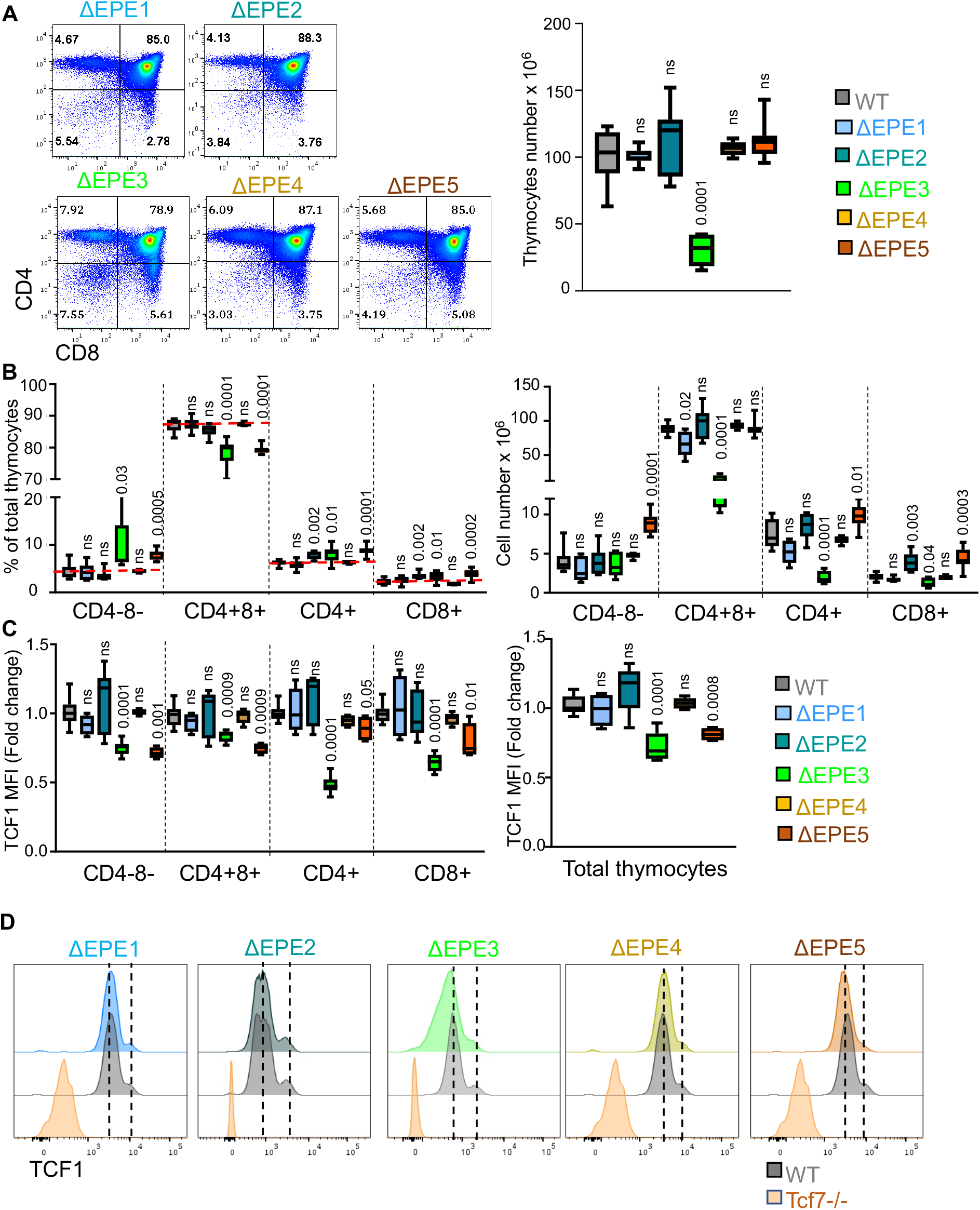
Impact of deletion of each EPE on αβ T cell development. **(A)** Representative histograms (left) of thymic subsets defined by CD4 and CD8 expression and absolute numbers (right) of thymocytes from thymi of ΔEPE1, ΔEPE2, ΔEPE3, ΔEPE4, and ΔEPE5 mice. Absolute number measurements of total thymocytes in the thymus of WT (n=10), ΔEPE1 (n=8), ΔEPE2 (n=6), ΔEPE3 (n=7), ΔEPE4 (n=8), and ΔEPE5 mice (n=10). **(B)** Graphical representation of the frequency and absolute cell numbers of conventional αβ T-cell subsets defined by CD4 and CD8 expression in thymus of WT (n=7), ΔEPE1 (n=8), ΔEPE2 (n=6), ΔEPE3 (n=7), ΔEPE4 (n=6), and ΔEPE5 mice (n=10). **(C)** Graphical representation of the fold-change in mean fluorescent intensity of TCF1 in total thymocytes and subsets from WT (n=6), EPE1 (n=4), EPE2 (n=6), EPE3 (n=8), EPE4 (n=5), and EPE5 mice (n=4). **(D)** Representative histograms of intracellular staining for TCF1 in total thymocytes of WT and homozygous ΔEPE1, ΔEPE2, ΔEPE3, ΔEPE4, ΔEPE5, and Δ*Tcf7*-*/-* mice. Dashed lines denote mean peak intensity for the two peaks in WT thymocytes.

**Supplementary Figure 4:**
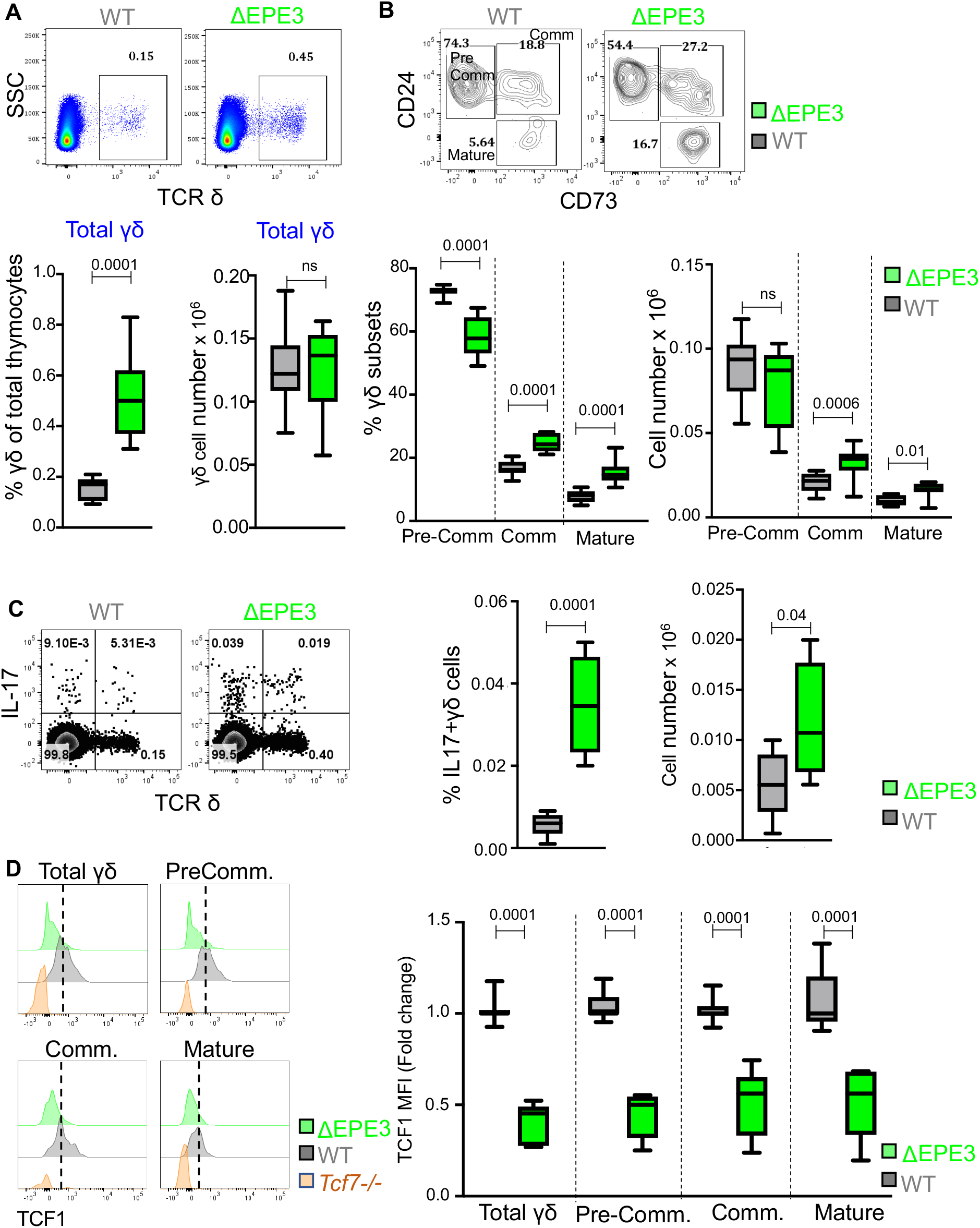
Impact of EPE3 ablation on TCF1 expression and development of γδ T-cells. (**A-C**) Flow cytometry analysis of the number and developmental progression of γδ T-cells by staining for thymocytes from WT and ΔEPE3 mice for surface expression of TCRδ, CD73 and CD24. **(A)** Graph represents the frequency (left) and absolute number (right) of γδ T-cells in the thymus of WT (n=13), and ΔEPE3 mice (n=11). (**B**) Two-color histograms of surface expression of CD24 and CD73 on TCR8+ cells (top) with developmental intermediates indicated (Precommitment - CD73-CD24+; Committed – CD73+CD24+; and Mature – CD73+CD24-) and a graphical representation of their frequency (left) and absolute number (right). WT (n=13), and ΔEPE3 mice (n=10). (**C**) Two-color histogram of flow cytometric analysis of IL17 production by TCR8+ thymocytes from WT, and ΔEPE3 mice, following activation with PMA and Ionomycin for 6 h. The frequency (left) and absolute number (right) of IL17 producing γδ T-cells in the thymi of WT (n=8), and ΔEPE3 mice (n=4) are depicted graphically. (**D**) Quantitation of TCF1 expression by intracellular flow cytometry in γδ T-cell developmental intermediates in the thymi of WT, ΔEPE3 and *Tcf7*-/- mice. Dashed lines denote mean peak intensity of TCF1 in WT thymocytes. Representative histograms (left) and fold-change in MFI of TCF1 staining in the indicated γδ T-cell subsets is shown. WT (n=7), and ΔEPE3 mice (n=5).

**Supplementary figure 5:**
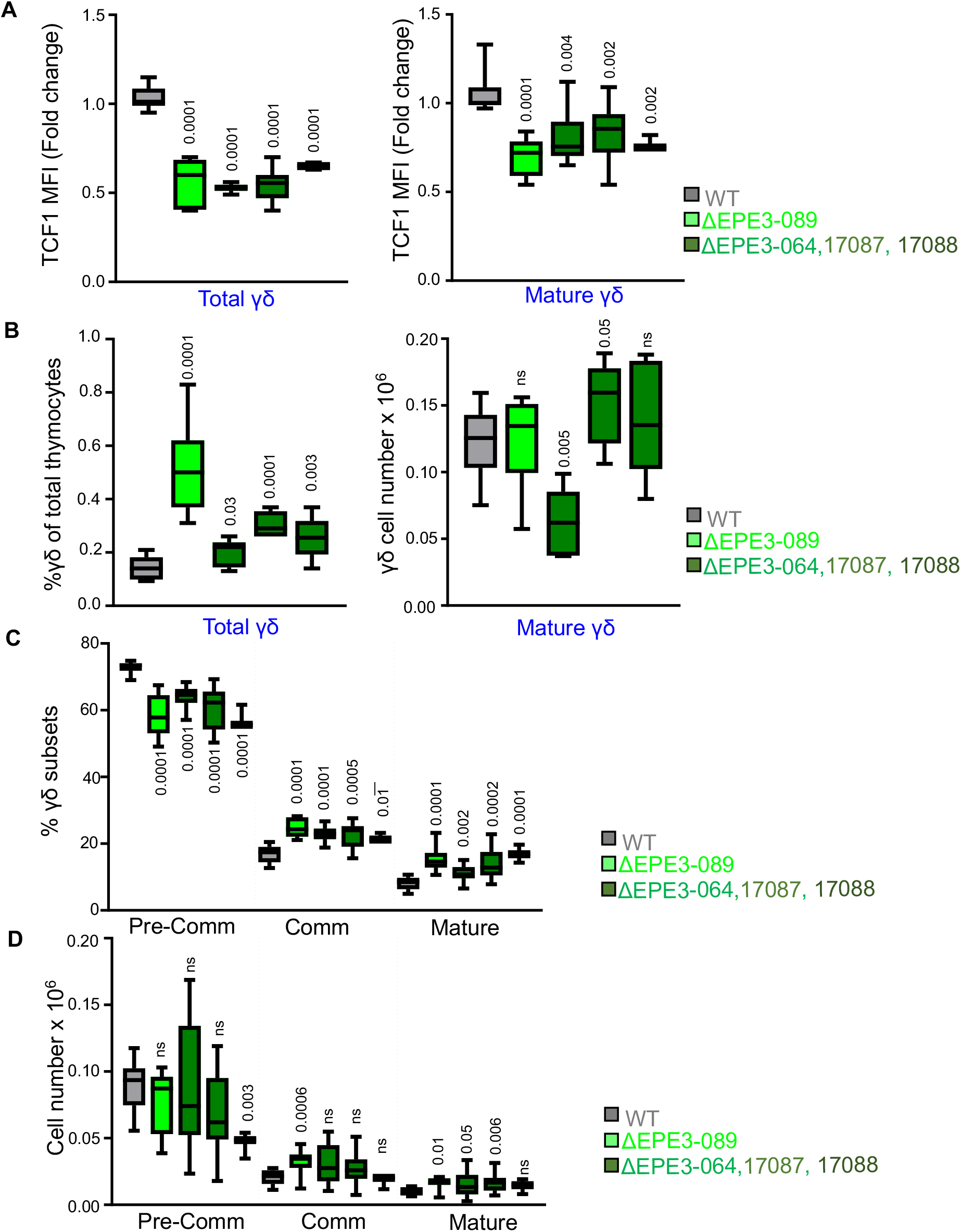
Impact of EPE3 ablation on TCF1 expression and development of γδ T-cells in other ΔEPE3 founder lines. **(A)** Graphical representation of the fold-change in MFI of TCF1 staining in total (left) and mature γδ T-cells (right) in WT (n=9), ΔEPE3-089 (n=5), ΔEPE3-064 (n=6), ΔEPE3-17087 (n=12), and ΔEPE3-17088 lines (n=3). **(B)** Graphical representation of the frequency (left) and absolute number (right) of total and mature γδ T cells in the thymi of WT (n=10), ΔEPE3-089 (n=6), ΔEPE3-064 (n=7), ΔEPE3-17087 (n=6), and ΔEPE3-17088 lines (n=6). **(C)** Graphical representation of the frequency of γδ T-cell subsets in the thymus of WT (n=10), ΔEPE3-089 (n=10), ΔEPE3-064 (n=16), ΔEPE3-17087 (n=17), and ΔEPE3-17088 lines (n=3). **(D)** Graphical representation of the absolute number of γδ T-cell subsets in WT (n=10), ΔEPE3- 089 (n=6), ΔEPE3-064 (n=7), ΔEPE3-17087 (n=6), and ΔEPE3-17088 lines (n=6).

**Supplementary figure 6:**
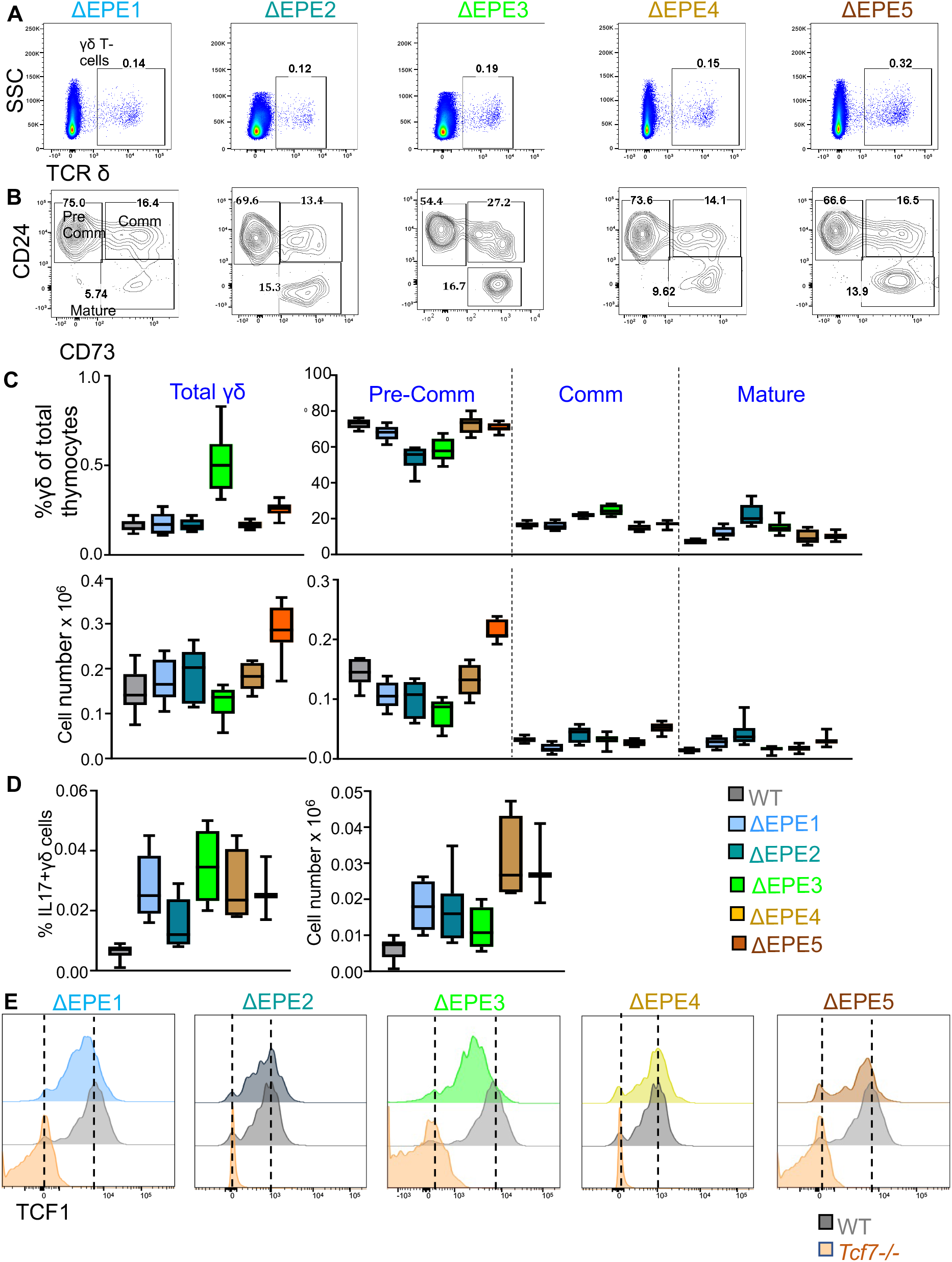
Effect of ablation of each EPE on development of γδ T-cells. **(A)** Representative histograms depicting the frequency of γδ T cells (TCRδ+ cells) in thymi from ΔEPE1, ΔEPE2, ΔEPE3, ΔEPE4, and ΔEPE5 mice. **(B)** Representative histograms of flow cytometric analysis of γδ T-cell developmental intermediates marked by staining thymocytes from ΔEPE1, ΔEPE2, ΔEPE3, ΔEPE4, and ΔEPE5 mice with antibodies to CD24 and CD73. **(C)** Graphical representation of the frequency (top) and number (bottom) of γδ T-cells and subsets in thymi of WT (n=8), ΔEPE1 (n=9), ΔEPE2 (n=6), ΔEPE3 (n=11), ΔEPE4 (n=8), and ΔEPE5 mice (n=10). **(D)** Graphical representation of the frequency (left) and absolute number (right) of IL17+ γδ T-cells in the thymus of WT (n=11), ΔEPE1 (n=6), ΔEPE2 (n=6), ΔEPE3 (n=6), ΔEPE4 (n=4), and ΔEPE5 mice (n=3). **(E)** Representative histograms of intracellular staining for TCF1 expression in γδ T-cells of ΔEPE1, ΔEPE2, ΔEPE3, ΔEPE4, and ΔEPE5. Dashed lines denote mean peak intensities of TCF1 in WT and *Tcf7*^-/-^ thymocytes.

**Supplementary figure 7:**
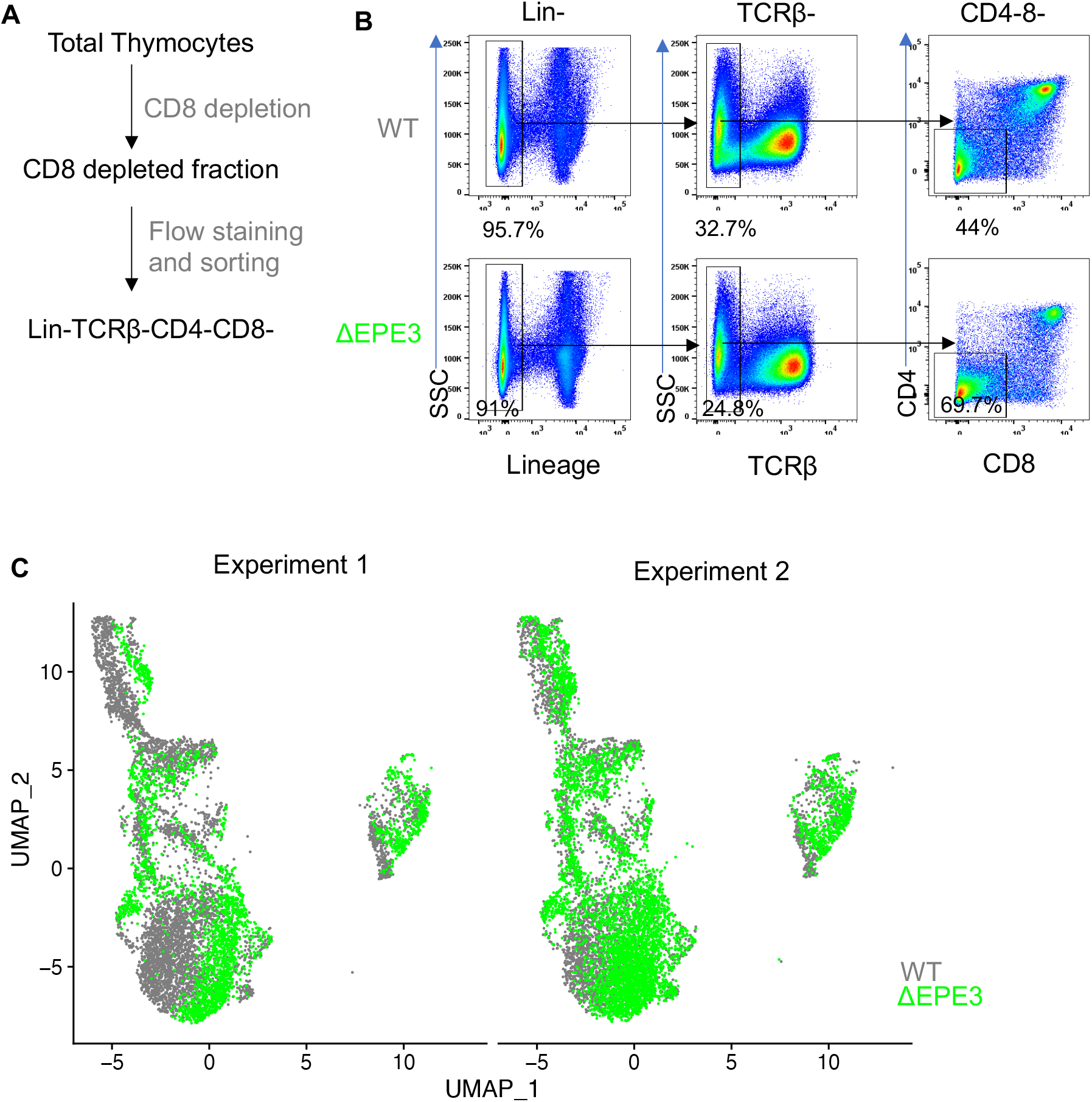
Sorting strategy of DN fraction for single cell RNA-seq. **(A)** Schematic representation of DN fraction sort. (Lineage- - defined as depleted with antibodies reactive to: TCRβ, CD4, and CD8, CD11c, CD11b, Gr1, CD19, B220, Ter119) **(B)** Flow cytometry profiles depicting the sorting strategy employed to isolate DN fractions from lineage depleted thymocytes from WT and ΔEPE3 mice, which were then used for scRNA-Seq analysis. **(C)** UMAP plots comparing subset distributions of thymocytes from WT and 1′EPE3 mice from two distinct experiments.

**Supplementary figure 8:**
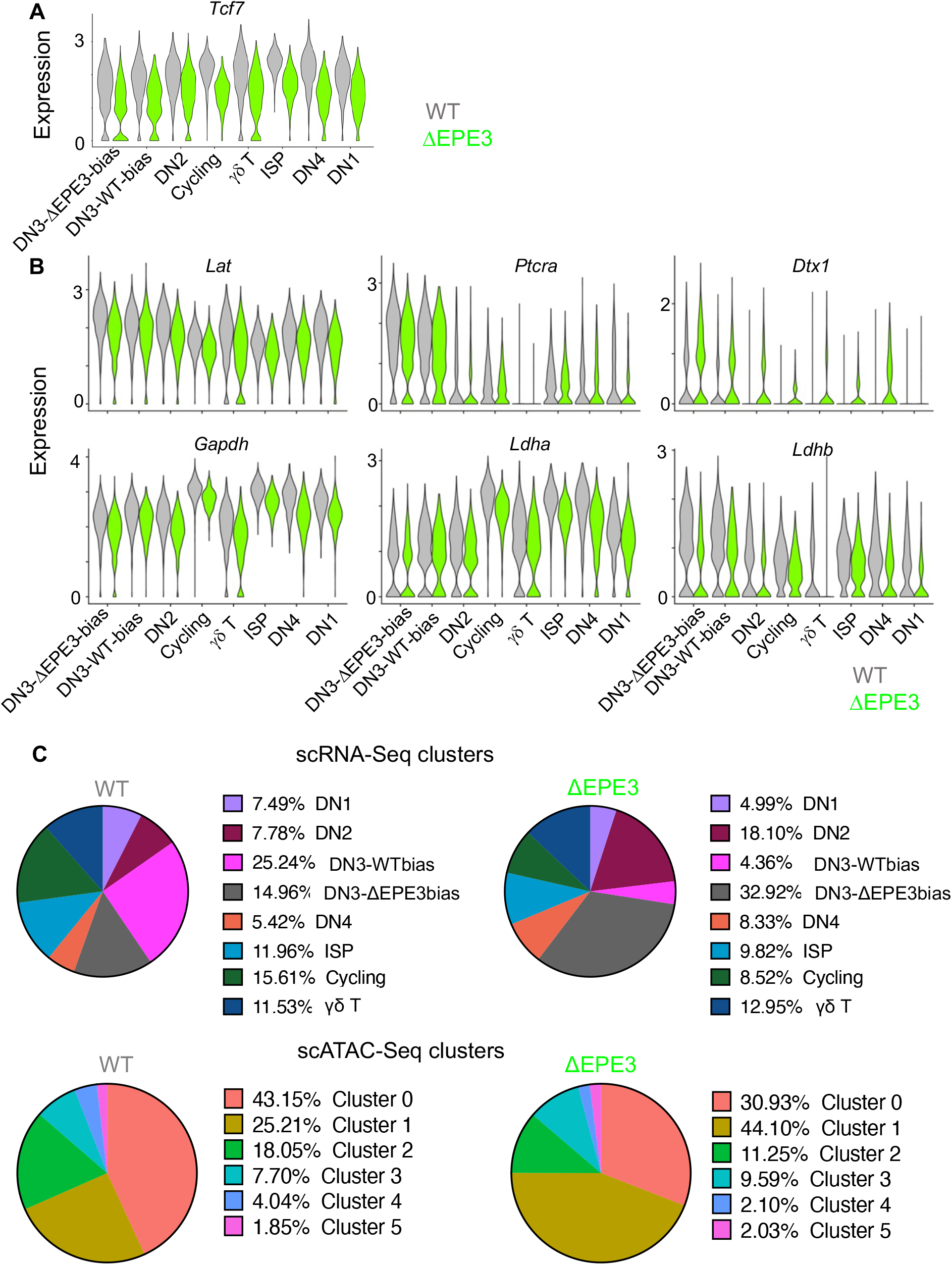
scRNA-Seq analysis of DN thymocytes from WT and 1′EPE3 mice. **(A)** Violin plots of *Tcf7* expression in all clusters identified in WT and 1′EPE3 mice. **(B)** Violin plots of differentially expressed genes of interest in all clusters identified in WT and 1′EPE3 mice. **(C)** Pie charts illustrating the abundance of DN subsets from WT and ΔEPE3 mice within the scRNA-Seq (top) and scATAC-Seq (bottom).

**Supplementary figure 9:**
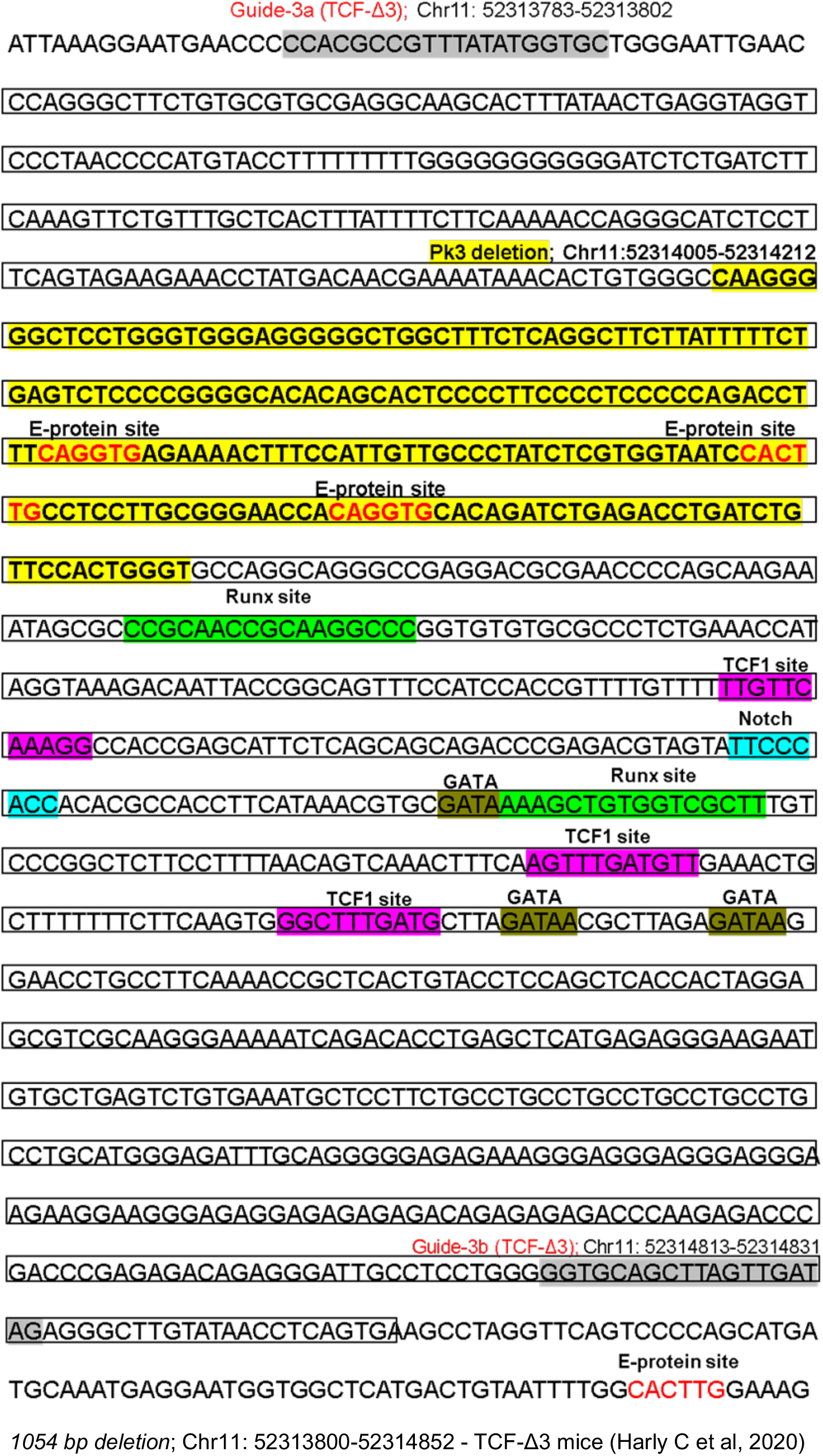
Motif analysis of binding sites surrounding EPE3. Sequence of chromosome 11 containing the previously published 1054 bp *Tcf*-*Δ3* deletion (Chr11:52313783- 52313802; termini marked by gray shading) ^21^. This genomic interval contains EPE3, which is shaded in yellow. Predicted transcription factor binding sites, including Notch, are indicated.

## Notes

### Competing Interest Statement

The authors have declared no competing interest.

